# Post-embryonic maturation of the *C. elegans* motor circuit

**DOI:** 10.1101/2022.01.24.477421

**Authors:** Ben Mulcahy, Daniel Witvliet, James Mitchell, Richard Schalek, Daniel Berger, Yuelong Wu, Doug Holmyard, Yangning Lu, Tosif Ahamed, Aravinthan Samuel, Andrew D Chisholm, Jeff Lichtman, Mei Zhen

## Abstract

During development, animals can maintain behavioral output even as the underlying circuits structurally remodel. After hatching, *C. elegans* undergoes substantial motor neuron expansion and synapse re-wiring while the animal continuously moves with an undulatory pattern. To understand how the circuit transitions from its juvenile to mature configuration without disrupting functional output, we reconstructed the *C. elegans* motor circuit by electron microscopy across larval development. We observed: 1) Embryonic motor neurons transiently interact with the developing post-embryonic motor neurons prior to remodeling of their juvenile wiring; 2) Post-embryonic neurons initiate synapse development with their future partners as their neurites navigate through the juvenile nerve cords; 3) Embryonic and post-embryonic neurons sequentially build structural machinery needed for the adult circuit before the embryonic neurons relinquish their roles to post-embryonic neurons; 4) This transition is repeated region by region along the body in an anterior to posterior sequence, following the birth order of post-embryonic neurons. Through this orchestrated, programmed and gradual rewiring, the motor circuit transforms from asymmetric to symmetric wiring. These maturation strategies support the continuous maintenance of motor patterns as the juvenile circuit develops into the adult configuration.

**Highlights:**

- Post-embryonic motor circuit maturation was reconstructed by synapse-resolution serial EM.
- Motor patterns are maintained as the circuit matures from asymmetric to symmetric configuration.
- Programmed rewiring gradually and sequentially transforms the circuit structure.
- Preparatory and communicative wiring-rewiring allows maturation without functional disruption.

## Introduction

The nervous system is built in embryos, but continues to expand and remodel after birth^1–3^. Postnatal growth is associated with changes in the body size, body plan, behavior, and habitat of the maturing animal. During postnatal development, neurons that are added or removed from a nervous system alter the wiring of functioning circuits. Animals require sensorimotor coordination for their survival. Critical circuitry must continuously maintain function even as it undergoes structural changes.

When the underlying circuits remodel, animals may update their behaviors or maintain their behaviors by compensating for structural changes. Before asking about the functional response to structural changes, we first need to characterize the structural changes across development. For most neural systems, this has not been described either with synapse resolution or at the level of full circuits.

The lineage of every cell of *C. elegans* has been mapped from the first cell division to adulthood^3,4^. Through lineage tracing^5^ and wiring reconstruction^6,7^, the *C. elegans* motor circuit was found to change dramatically during post-embryonic stages^5,6^. After hatching from embryos, the number of motor neurons that contribute to body bending expands from 22 to 75, and the number of neuron classes increases from 3 to 7^5^. One class, the embryonically born GABAergic neurons (DD), reverses neurite polarity: their dendrites in the first larval (L1) stage become the axons in the adult, and their axons become the dendrites^6,8^. Both axons and dendrites have different synaptic partners between the L1 and adult stages^6,9^.

Since the discovery of DD remodeling, the *C. elegans* field has turned to identifying its molecular determinants^10–13^. In these studies, DD motor neurons were labelled with fluorescent synaptic markers, and candidate remodeling genes were identified based on their effects on marker expression patterns. However, marker studies alone do not offer a full account of the cellular processes that convert the L1 motor circuit to its adult form. Without knowing precisely how structural changes occur in developing larvae, we cannot confidently place identified molecular pathways in context of cellular events.

Our study sought to fill this gap by longitudinal, synapse-resolution serial-section electron microscopy (EM) reconstruction of the developing *C. elegans* motor circuits. We fully reconstructed the body nerve cords of three larvae aged before, during, and immediately after the birth of post-embryonic motor neurons, and partially reconstructed two larvae at later time points. We observed a continuous and gradual transition from the juvenile circuit into the adult form: across early larval stages and along the body axis, motor neurons sequentially build structures for their adult stage configuration before dismantling their juvenile connections.

A full account of structural maturation is critical to resolve a standing mystery of postnatal development: despite the substantial changes in motor circuit anatomy, *C. elegans* performs locomotion with similar ventral-dorsal bending waves across life stages^14,15^. How might post-embryonic expansion and remodeling occur without disrupting mobility or altering the motor pattern?

In another study, we explain how the newborn L1 larvae and adults generate similar body bends using different mechanisms^16^. Unlike the adult motor circuit, which is wired into two sub-circuits, each dedicated to either dorsal or ventral bending, a newborn L1 motor circuit is structurally wired to drive dorsal bending only. L1’s ventral bending takes place through the extrasynaptic activation of ventral muscles and anti-phasic entrainment with dorsal bending^16^. With two sets of mechanisms in place to produce the same behavioral output, a gradual transition from the juvenile to adult configuration allows *C. elegans* to maintain a uniform motor pattern throughout postnatal development.

In this study, we describe the temporal and spatial events of anatomical changes of the motor circuit, spanning post-embryonic neurogenesis, neurite migration, synapse formation, and synapse disassembly. We highlight three strategies by which the juvenile circuit transits to its adult form without functional interruption. One is the sequential remodeling of adjacent motifs of the motor circuit along the body. When the motor circuit undergoes remodeling one segment at a time, the similar output of juvenile and adult configurations ensures continuity of bending along the body. Another is the preparatory remodeling, where motor neurons initiate structural changes in anticipation of the adult-stage configuration before the juvenile configuration is dismantled. Preparatory remodeling allows a seamless transition with flexible timing along the body axis. Last is the communicative remodeling, where motor neurons that swap wiring partners transiently interact prior to the exchange taking place. Physical interactions work in conjunction with intrinsic programs to coordinate wiring replacement.

A detailed description of these morphological events that are necessary to facilitate downstream mechanistic studies is included in Results. Readers can also proceed directly to Discussion where historic contexts, general conclusions, and broad implications are summarized.

## Results

### Post-embryonic motor circuit development establishes wiring symmetry

During post-embryonic development, *C. elegans* distributes 53 new body-bending motor neurons along the L1 larva’s ventral nerve cord^5^ (illustrated in Fig. 1A). Motor neuron wiring between the newborn L1 and adult motor circuits also differs substantially^6,7,16^ (illustrated in Fig. 1B).

**Figure 1.**
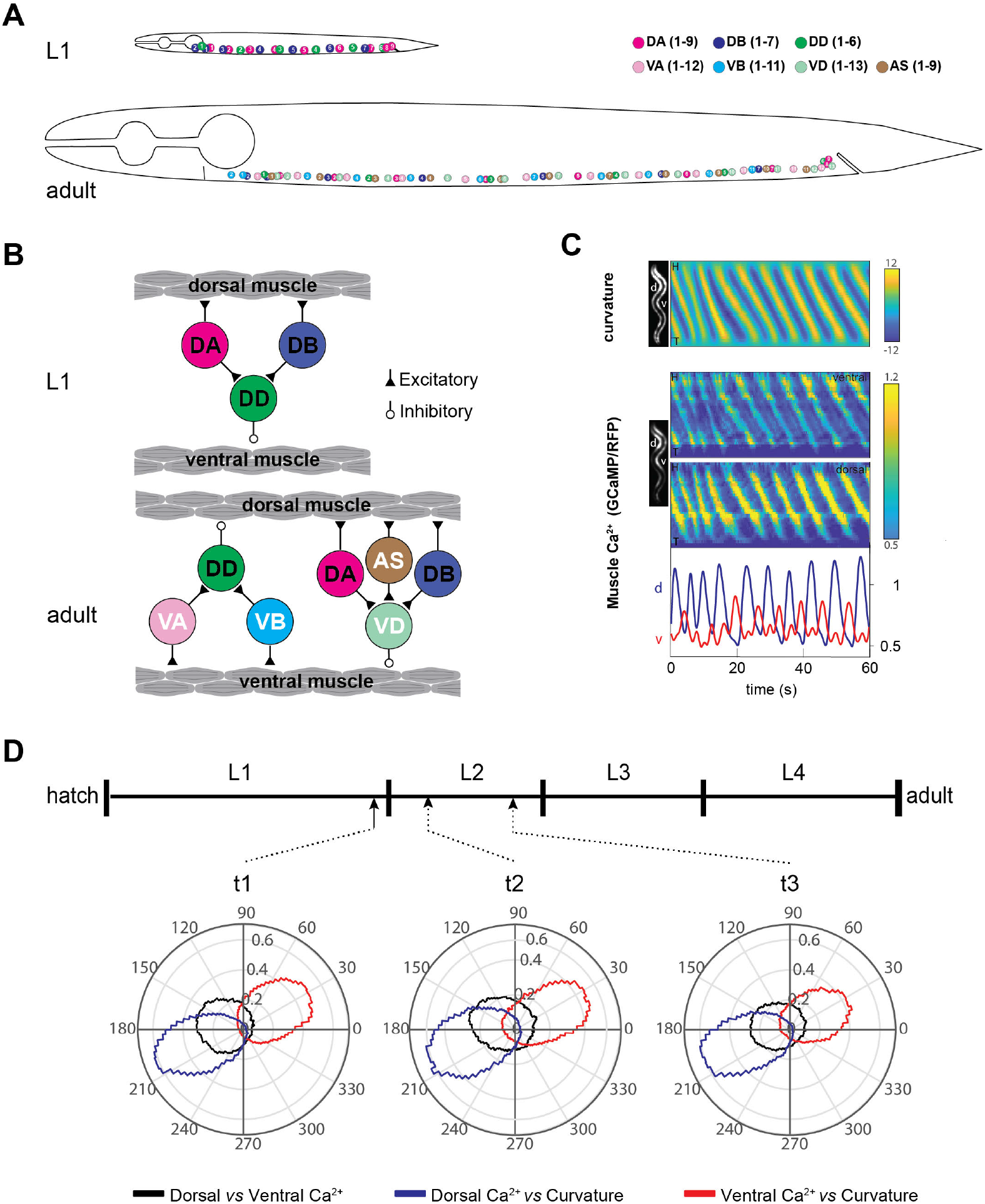
*C. elegans* larvae maintain coordinated motor output as the motor circuit matures. (A) Body motor neurons that belong to different classes are distributed across the ventral nerve cords of newborn L1 larva and adult animals (body size not to scale; VC motor neurons not shown). (B) Schematic wiring diagram between body motor neurons and body wall muscles in L1 and adult animals. (C) Representative kymographs of a freely crawling larva (t2 in D) expressing GCaMP3::RFP in body wall muscles. Top: curvature along the body over time. Bending propagates from head (segment 1) to tail (segment 30), representing a period of forward undulation. Middle: calcium signals, represented by GCaMP3/RFP ratio, in ventral and dorsal muscles along the body. Bottom: example calcium signals of the ventral (blue) and dorsal (red) muscles, extracted in the mid-body (segment 18). (D) Polar histograms quantifying phase differences along the body (segments 11-30) between dorsal muscle activity and curvature (blue), ventral muscle activity and curvature (red), and dorsal and ventral muscle activity (black), at the onset, middle and end of the juvenile-to-mature motor circuit transition (n = 10-12 animals).

A newly hatched L1 larva has 22 motor neurons that belong to three classes^6^. One set of inhibitory motor neurons (iMNs), the GABAergic DD neurons, relax ventral muscles, while two sets of excitatory motor neurons (eMNs), the cholinergic DA and DB neurons, contract dorsal muscles. These eMNs and iMNs are wired in repeating motifs along the anterior-posterior axis, forming a circuit for dorsal bending that coordinates contraction of dorsal muscles with relaxation of opposing ventral muscles^6,16^ (Fig. 1B, upper panel). Lacking a separate circuit to directly drive ventral bending, the L1 larva entrains ventral bending to the activity of the dorsal bending circuit^16^.

Adult *C. elegans* adds four new classes of motor neurons for body-bending^9^ and also adds a ventral bending circuit^9^ (illustrated in Fig. 1B, lower panel). Three new sets of cholinergic eMNs contract either ventral muscles (VA and VB eMNs) or dorsal muscles (AS eMN). One new set of GABAergic iMNs, the VD neurons, relax ventral muscles. The embryonic set of iMNs, DD, is remodeled to relax dorsal muscles. iMNs and eMNs are wired to form repeating motifs of two sub-circuits that directly drive bending. In each sub-circuit, eMNs that contract muscles synapse onto iMNs that relax muscles on the opposite side, thereby making either dorsal or ventral bends^9^.

Thus, post-embryonic development adds both structural and functional symmetry to create the adult motor circuit. This requires the L1 motor circuit to integrate new classes of motor neurons and reconfigure the wiring of all embryonic motor neurons.

### Post-embryonic motor circuit development does not disrupt or alter motor output

We asked whether post-embryonic remodeling causes any interruption of motor patterns. Post-embryonic cell divisions give rise to motor neurons between the mid-L1 and early-L2 stages^5^. When these motor neurons begin to integrate into the motor circuit was unknown. We thus quantified undulatory movements across age, covering periods before, during, and after motor neuron birth. We observed nearly identical undulatory patterns at all ages, where symmetric dorsal-ventral bending waves traveled from head to tail (Fig 1C, D; Fig 1 – Supp. 1).

All bending waves were strongly correlated with muscle activities. By imaging calcium dynamics in body wall muscles of crawling larvae, we found that bending wave propagation is spatially and temporally correlated with waves of muscle activity that propagate along the body in the same way at all ages (Fig 1C, D). Thus, during post-embryonic development, muscle activity and locomotion patterns continue without disruption or alteration while the motor circuit remodels.

### Deducing post-embryonic motor circuit development by high-resolution EM reconstruction of six larvae

How does the L1 motor circuit make an uninterrupted transition into its adult configuration? Remodeling of L1’s dorsal bending circuit and integration of the ventral bending circuit must be orchestrated across motor neurons and along the body. To understand this orchestration, we performed longitudinal synapse-resolution serial-section EM reconstruction of the developing motor circuit. Previously, using differential interference contrast microscopy, Sulston deduced the birth order of all motor neurons by tracking the cell nuclei^5^. Starting at the mid-L1 stage and ending at the early second larva (L2) stage, thirteen precursor (P) cells sequentially migrate into the ventral nerve cord, divide consecutively, and give rise to new motor neurons in an anterior-to-posterior temporal order (illustrated in Fig. 2A). When these motor neurons start to extend neurites and to make synapses was unknown. But this stereotypy, combined with other lineage landmarks^5^, allowed us to identify every cell in our EM datasets and to assign ages to our six samples (Supplemental Table 1).

**Figure 2.**
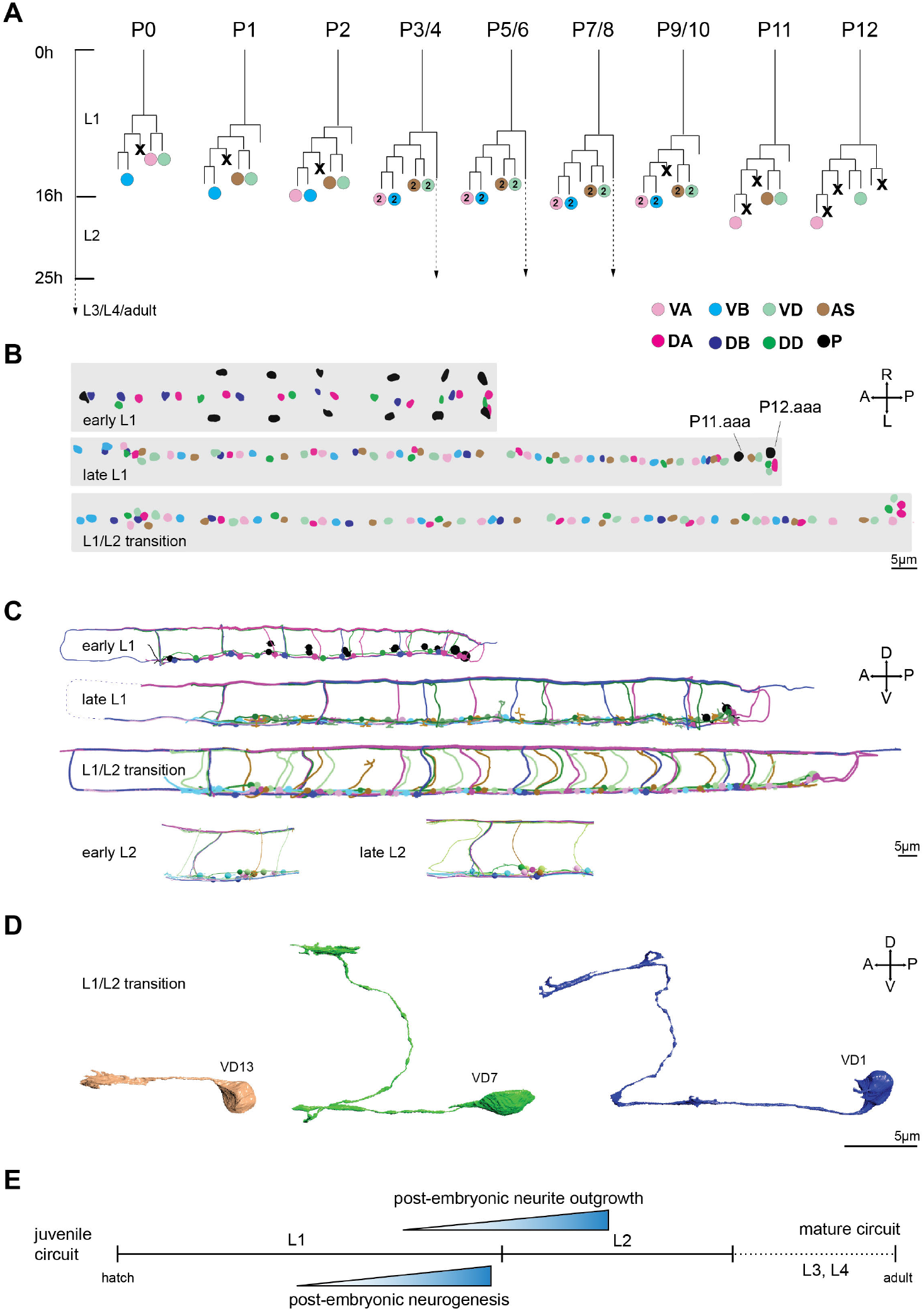
Post-embryonic motor neuron birth and growth follows a stereotyped process. (A) The lineage tree of post-embryonic motor neurons. Thirteen precursor cells (P0-12) give birth to 5 new classes of motor neurons by several rounds of division. Each horizontal line denotes one stereotyped cell division event that occurs between the late first (L1) and early second (L2) larva stage. Only neuronal progeny are denoted, and the developmental timing is based on animals raised at 20°C. Circles: neurons colored coded by classes; X: apoptosis. Adapted from^5^. (B) A dorsal view of nuclei of differentiated motor neurons and motor neuron precursors along the ventral nerve cord, reconstructed from serial electron micrographs of whole larvae at early L1 (1hr), late L1 (15hr) and the L1/L2 transition (16hr). Precursors P1-12 reside laterally immediately after birth (1hr). They sequentially migrate into the ventral nerve cord and divide following an anterior to posterior order. At late L1 (15hr), P0-10 have given birth to motor neurons whereas P11 and P12 have just entered the ventral nerve cord. (C) Skeletal reconstruction of embryonic and post-embryonic motor neurons at respective developmental ages. Circles denote neuron soma scaled to their nuclei size; lines denote neurites. At late L1 (15hr) and the L1/L2 transition (16h), post-embryonic motor neurons are at various stages of neurite outgrowth, roughly following the birth order (anterior-to-posterior). (D) Examples of stereotyped morphology of post-embryonic motor neurons undergoing neurite outgrowth at the L1/L2 transition. Reconstruction of VD class motor neurons during three phases of neurite outgrowth. *(Left)* During early stages of outgrowth, a nascent neurite adopts an amorphous, large profile that fills space and wraps other neurites in the ventral nerve cord. *(Middle)* During commissural outgrowth, sheet-like and branched growth cones lead dorsal migration, interacting with epidermis, excretory canal and sub-lateral nerve cords on their paths. *(Right)* After reaching the dorsal nerve cord, the process expands into an amorphous large profile, filling space and wrapping around other neurites as it expands anteriorly and posteriorly along the dorsal nerve cord. (E) Schematics summarizing the temporal order of post-embryonic neurogenesis and neurite outgrowth in the ventral nerve cord.

We began with fully reconstructing the body of three larvae aged between the L1 and L2 larval stages (Fig. 2B). These reconstructions captured motor circuit development before (early L1), during (late L1), and after (L1/L2 transition) the birth of most post-embryonic motor neurons (Fig. 2 – Supp. 1). A second partially reconstructed young L1 larva prior to post-embryonic neurogenesis recapitulates the early L1 larva’s pattern^16^. To capture events towards completion of remodeling, we further reconstructed the anterior nerve cords of two older larvae at early and late phases of the L2 stage (Fig. 2C).

In fully reconstructed larvae, because members of the same neuron class are born sequentially, we can deduce the developmental progression of each class by comparing its members along the body. Because each precursor cell contributes one member for each new motor neuron class (Fig. 2A), we can also deduce the developmental progression of circuit reconfiguration by comparing the wiring between adjacent motifs along the body.

By comparing motor neurons of the same class along the anterior-posterior axis and comparing the same motor neurons at different developmental age, we were able to deduce a sequence of cellular events that occur in a stereotyped and iterated manner. Our observations reveal an organized orchestration of wiring reconfiguration that leads to the adult-like wiring pattern by the end of L2 stage across the body.

Below we describe remodeling through three parallel events: neurite development of nerve cords (Fig. 2, Fig. 3); remodeling of the dorsal bending circuit (Fig. 4); building of the ventral bending circuit (Fig. 5). We note that these are mutually-dependent organic events. Our categorization represents an arbitrary demarcation, adopted for simplicity of presentation.

**Figure 3.**
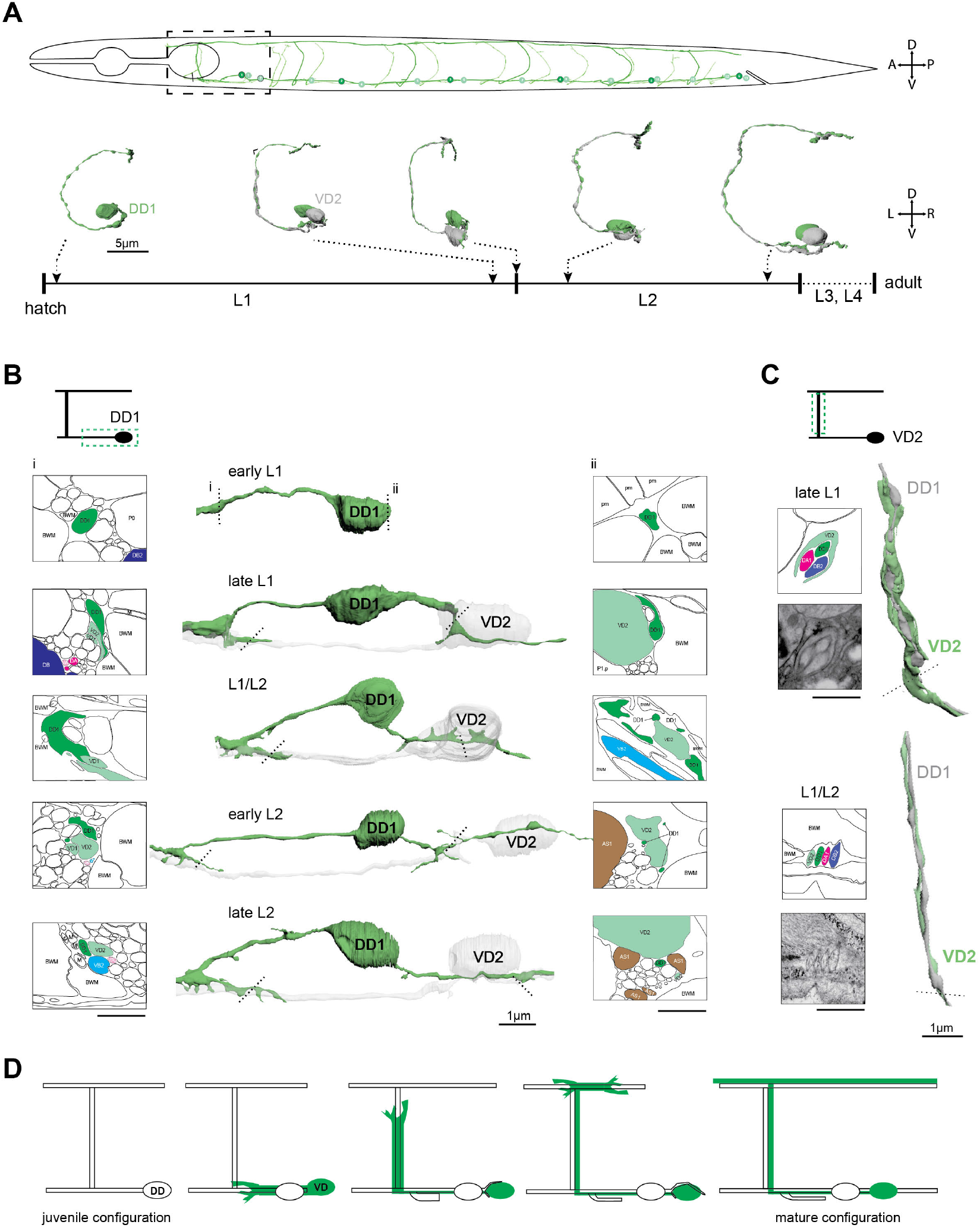
Neurites of embryonic and post-embryonic iMNs transiently interact during post-embryonic development. (A) (Upper panel) Skeleton segmentation of iMNs of the DD and VD classes at the L1/L2 transition superimposed along the body axis of a worm. DD1 and VD2 were chosen to represent their respective classes in additional reconstructions at later time points. (Lower panel) A tilted posterior-to-anterior view of the volumetrically reconstructed DD1/VD2 iMN pair at five developmental time-points. Hours denote the developmental age of larvae raised at 20°C in reference to anatomical landmarks. (B) (Center) Enlarged views of migrating DD1 and VD2 neurites in the ventral nerve cords at different developmental ages, with morphologies at indicated positions shown in illustrated EM cross-sections. Left and right panels are membrane outlines of the EM cross-section profiles marked by dashed lines at each stage. (C) (Right) VD2 commissure outgrowth proceeds through wrapping around an existing fascicle containing DD1, DA1, and DB2 commissures (late L1), then unwraps to run in parallel with other neurites in the commissure bundle (L1/L2 transition). (Left) Membrane outlines of the EM cross-section profiles marked by dashed lines. (D) Schematics summarizing transient interactions between embryonic and post-embryonic iMN neurites during post-embryonic development.

**Figure 4.**
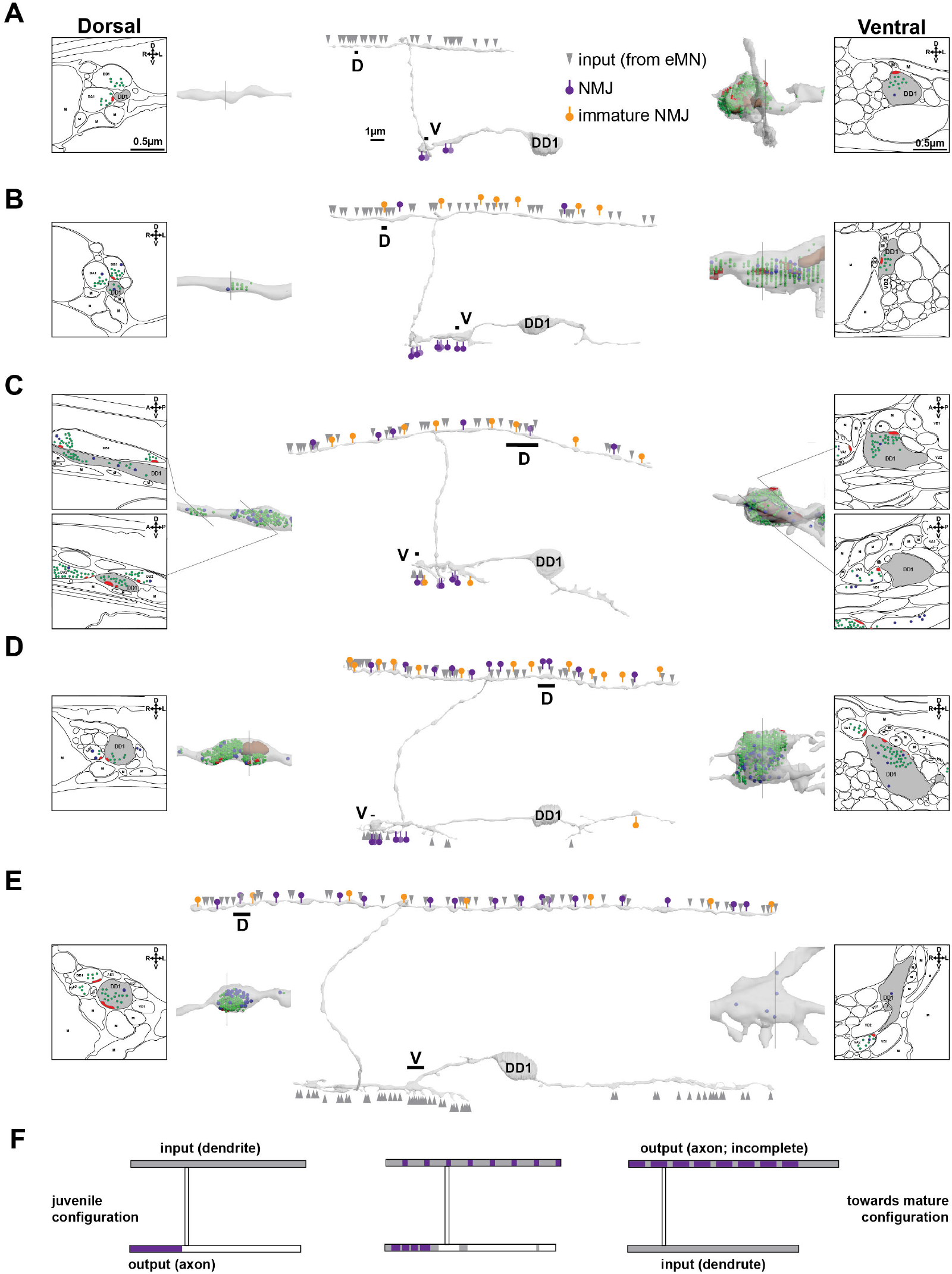
Synaptic rewiring of embryonic iMNs rebuilds the dorsal bending circuit. (A-E) Volumetric reconstruction of DD1 across developmental ages, from early L1 (A), late L1 (B), the L1/L2 transition (C), early L2 (D), to late L2 (E). Center panels are full models of DD1 soma and processes; enlarged views of a region of the dorsal and ventral processes are shown in the left and right panels, respectively. Example EM cross-sections (labelled by dashed lines) are shown with traced membrane outlines. In center panels, grey arrowheads represent synaptic inputs to DD1; purple arrows with closed circles denote presynaptic termini of DD1, which are NMJs or dyadic synapses with muscles as co-postsynaptic partner. Orange arrows with closed circles denote immature presynaptic termini of DD1, consisting of swellings with synaptic vesicles, but no discernible presynaptic dense projections. In the left and right panels, synaptic vesicles (green), dense core vesicles (blue), presynaptic dense projections (red), and mitochondria (brown) are shown. (F) Schematics summarizing sequential events of DD rewiring with their pre-and post-synaptic partners during post-embryonic development.

**Figure 5.**
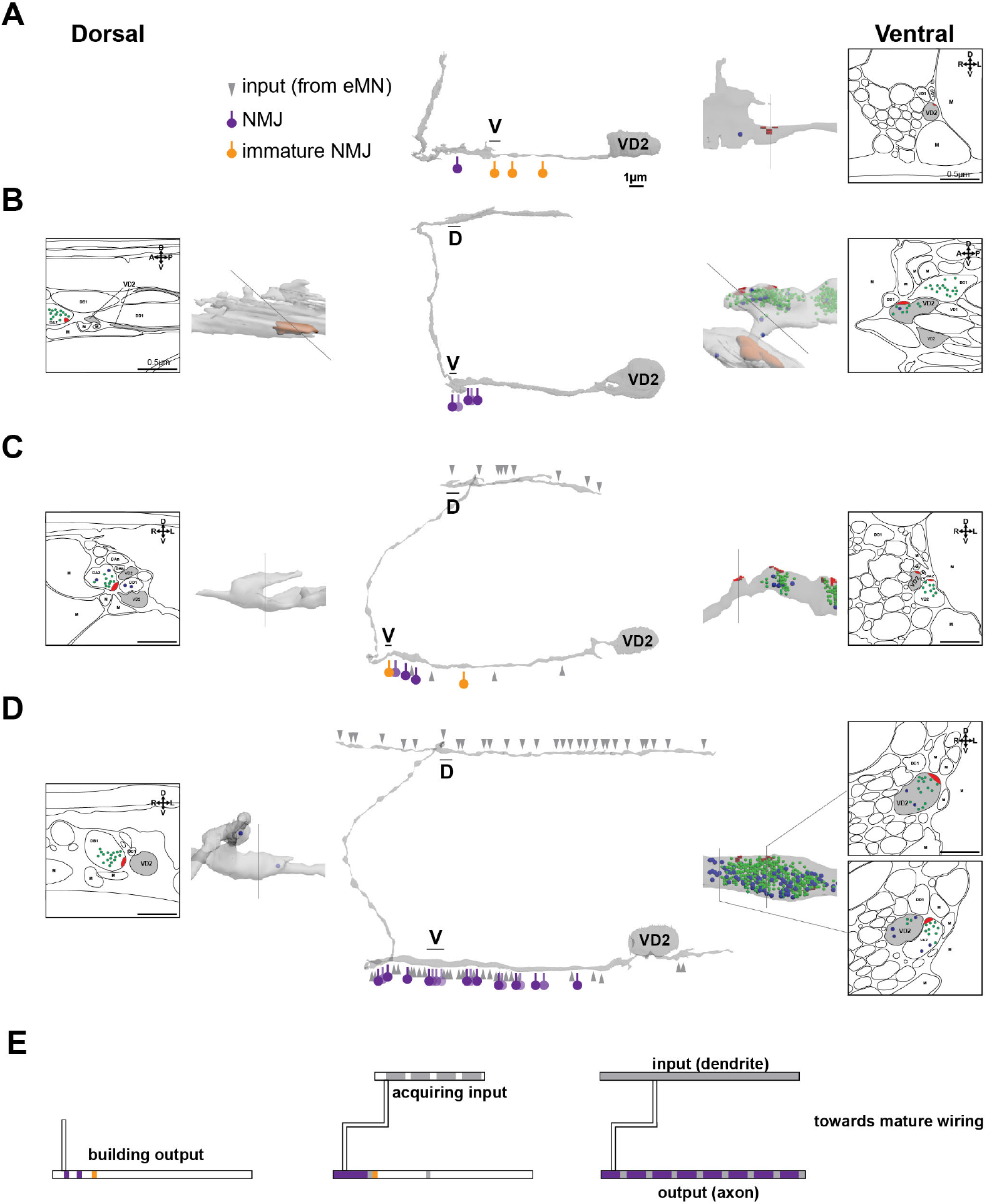
Post-embryonic iMNs take over embryonic iMN’s synaptic partners to build the adult ventral bending circuit. (A-E) Volumetric reconstruction of VD2 across developmental ages, from early L1 (A), late L1 (B), the L1/L2 transition (C), early L2 (D), to late L2 (E). Center panels are full models of VD2 soma and processes; enlarged views of a region of the dorsal and ventral processes are shown in the left and right panels, respectively. Example EM cross-sections (at dashed lines) are shown with traced membrane outlines. In center panels, grey arrowheads represent synaptic inputs to VD2; purple arrows with closed circles denote presynaptic termini of VD2, which are NMJs or dyadic synapses with muscles as co-postsynaptic partner. Orange arrows with closed circles denote immature presynaptic termini of VD2, consisting of swellings with presynaptic dense projections but no discernible cloud of synaptic vesicles. In the left and right panels, synaptic vesicles (green), dense core vesicles (blue), presynaptic dense projections (red) and mitochondria (brown) are shown. (F) Schematics summarizing sequential establishment of VD’s pre-and post-synaptic partners during post-embryonic development.

#### Post-embryonic neurite development of the dorsal and ventral nerve cords

After birth, the embryonic DD iMNs continue to extend neurites along both the dorsal and ventral nerve cords. In the newborn L1, DD neurites do not yet fully cover both nerve cords, but each neuron makes at least one physical contact with the neurite of a neighboring DD neuron. By late L1, all neurites are fully extended, tiling both dorsal and ventral cords with gap junctions (Fig. 3 – Supp. 1A(d)) between neighboring neurites (Fig. 2 – Supp. 3B).

In the newborn L1, the nuclei of precursors for motor neurons, the P cells, have not started their migration into the ventral nerve cord. However, all P cells have already extended cellular processes that line the ventral nerve cord (Fig. 2 – Supp. 2A). These processes guide the descent of nuclei into the cord, which occurs later in an anterior-to-posterior temporal order^3,4^. After this migration, each P cell gives rise to one member of each new neuron class (Fig. 2A, Fig. 2 – Supp. 1) with some descendants undergoing apoptosis (Fig. 2 – Supp. 2B).

After their birth, all four classes of post-embryonic motor neurons begin neurite extension along the ventral nerve cord. Two classes (VA and VB) develop only ventral processes (Fig. 2 – Supp. 3B, upper panels), while two classes (AS and VD) further extend commissures that project dorsally and run along the dorsal nerve cord (Fig. 2 – Supp. 3B, lower panels).

In this developing circuit, all neurites follow a similar migration process (Fig. 2 – Supp. 4, Supplemental data sheets 1-5). They extend with a sheet-like leading edge reminiscent of the axonal/dendritic growth cones^17,18^, similar to those observed for motor neurons during embryogenesis^19^. They wrap around existing tissues, detach from them, and leave behind a mature, tube-like nerve fiber that runs parallel to the initial support (Fig. 3C).

Some tissues exhibit interactions with all members of the same neuron class, making them potential guideposts. For dorsal-extending commissures, they include the sub-lateral nerve cords, the excretory canal, and neurites of embryonically born motor neurons (Fig. 3 – Supp. 2, Supplemental data sheets 1-3). Along the dorsal and ventral nerve cords, neurites wrap or attach to epidermis or neurites of embryonic motor neurons (Fig. 3 – Supp. 2, Supplemental data sheets 1-5).

#### Orchestrated establishment of new dorsal bending and ventral bending motor sub-circuits

Full maturation from the L1 to adult wiring configuration requires all embryonic motor neurons to establish new synaptic connections with post-embryonic motor neurons. These connections add a new circuit for ventral bending and reconstruct the circuit for dorsal bending.

To form the ventral bending circuit, embryonic iMNs (DD neurons) must disconnect from their original presynaptic partners, the embryonic eMNs (DA and DB neurons), and build new input connections from the post-embryonic eMNs (VA and VB neurons). The DD iMNs must also switch their postsynaptic partners from the ventral to dorsal muscles. VA and VB neurons form excitatory neuromuscular junctions (NMJs) to ventral muscles; their activation simultaneously activates DD to relax dorsal muscles, producing a ventral bend (illustrated in Fig. 1B, left side of the adult panel).

DD iMNs relinquish their role in dorsal bending to the post-embryonic VD iMNs. To rebuild the dorsal bending circuit, the VD iMNs must establish input connections from eMNs that excite dorsal muscles, the embryonic DA and DB neurons and the post-embryonic AS neurons. The VD iMNs must also establish inhibitory NMJs to ventral muscles. When DA, DB, and AS excite dorsal muscles, they simultaneously activate VD to relax ventral muscles, producing a dorsal bend (illustrated in Fig 1B, right side of the adult panel).

We found that maturation events are also iterated from anterior to posterior along the motor circuit. We combined reconstruction of three full circuits and three partial circuits to deduce the sequence of this process (Fig. 2C). In the anterior body, the embryonic iMN (DD1) and post-embryonic iMN (VD2) lead the wiring transition from the juvenile to mature configuration (Fig. 3A). Three morphological events are critical to the wiring reconstruction (below).

#### Embryonic and post-embryonic iMNs transiently interact, coinciding with the initiation of wiring transition

Developing DD1 and VD2 neurites interact transiently, which might facilitate the wiring transition (Fig. 3, Fig. 3 – Supp. 1B(*e*), C(*g*)). In the newborn L1, DD1 has an anteriorly projecting axon in the ventral cord and an anteriorly projecting dendrite in the dorsal cord. Neither process is branched (Fig. 3C). By the end of the L1 stage, the DD1 axon has acquired two posterior-projecting branches (Fig. 3B). One branch projects from the soma and finishes tiling with the neighboring DD2 axon. During its growth, it meets the VD2 soma in the L1 stage, wraps around it, and unwraps from it by the end of L2 stage. The other branch projects posteriorly from the proximal end of the axon, makes contact, then runs alongside the anterior-projecting neurite of VD2.

VD2’s commissure wraps around a fascicle of commissures of embryonic motor neurons, including DD1, as it climbs dorsally (Fig. 3C). When it reaches the dorsal cord, it bifurcates; both posterior and anterior projections extend while wrapping neurites of DD1 and another neuron, RID (Fig. 3 – Supp. 2A(*c*), B(*f*)).

The transient physical interactions between DD1 and VD2 neurites may provide mutual cues for wiring replacement, because the VD2 neurite adopts appropriate positions in the nerve cords as it extends. It develops its ventral presynaptic termini in the area initially inhabited by DD NMJs and takes over embryonic eMN input as it grows along the dorsal nerve cord (see below).

#### Embryonic iMNs change axon-dendrite identity, building the ventral bending circuit with new pre-and post-synaptic partners

In the newborn L1, DD’s ventral process is the axon, which makes inhibitory NMJs to ventral muscles. DD’s dorsal process is the dendrite, which receives inputs from embryonic DA and DB eMNs. In adults, these processes reverse their identities: the ventral process becomes the dendrite, receiving inputs from post-embryonic VA and VB eMNs, and the dorsal process becomes the axon, making inhibitory NMJs to dorsal muscles (Fig. 4 – Supp. 1).

Thus, in the newborn L1, DD iMNs are part of the dorsal bending circuit, facilitating relaxation of ventral muscles when dorsal muscles are activated. In the adult, the DD iMNs are part of the ventral bending circuit, facilitating relaxation of dorsal muscles when ventral muscles are activated. The striking feature of this identity reversal is that it does not involve breakdown of embryonic neurites. Instead, existing neurites undergo re-assignment of axonal and dendritic identities during their post-embryonic extension.

At birth, DD dendrites are post-synaptic to the axons of embryonic DA and DB eMNs in the dorsal nerve cord (Fig. 4A, Dorsal; Fig. 4 – Supp. 2A(*d, e*)). Beginning in late L1, the DD dendrite gradually acquires presynaptic morphology that eventually transforms the dorsal process into the axon. Nascent presynaptic termini appear along this process, first as a few small swellings with a few vesicles (Fig. 4B, Dorsal; Fig. 4 – Supp. 2B(*g*), C(*i*)). As the larva grows, these swellings acquire presynaptic dense projections, and increase in size and number. Almost every emerging presynaptic terminal apposes a muscle arm, establishing NMJs with the dorsal muscles (Fig. 4C-E, Dorsal; Fig. 4 – Supp. 2B(*f*), C(*h*)).

By late L1, the DD axon begins to exhibit postsynaptic morphology that transforms the ventral process into the dendrite (Fig. 4C, Ventral). It starts by sending spine-like structures towards swellings on the developing neurites of post-embryonic VA and VB eMNs (Fig. 3B (left panels); Fig. 3 – Supp. 2C(*f*)). By the L1/L2 transition, its anterior region apposes maturing presynaptic termini from these eMNS (Fig. 4C). As the larva grows, this ventral process receives more presynaptic inputs and develops more dendritic spines as it elongates (Fig. 4D-E).

Before remodeling is complete, both the dorsal and ventral neurites have a mixed identity, simultaneously exhibiting features of axons and dendrites. During the first larval stage, the anterior region of DD1 increases the number of NMJs it makes to ventral muscles, to support its role as a juvenile axon (Fig. 4A-D, Ventral). As the dorsal process develops presynaptic termini to the dorsal muscles, the same neurite retains its presynaptic inputs from the embryonic DA and DB eMNs to support its role as the juvenile dendrite (Fig. 4A-D, Dorsal). We only observed the disappearance of NMJs from the ventral process after the appearance of many mature-looking NMJs along the dorsal process (Fig. 4E).

In summary, during post-embryonic development, neurites of embryonic DD iMNs undergo a gradual switch of axon-dendrite identity as they establish new inputs and new outputs. Both neurites exhibit mixed axon-dendrite morphologies during this transition. The juvenile connectivity is only dismantled after the adult-stage connectivity is fully formed (Fig. 4F).

#### Post-embryonic iMNs rebuild the dorsal bending circuit, replacing embryonic iMNs as their neurites extend

As the embryonic DD iMNs establish new wiring, the post-embryonic VD iMNs take over their former pre-and post-synaptic partners to make the new dorsal bending circuit (Fig. 5 – Supp. 1). The striking feature of this process is that post-embryonic neurons build synapses during neurite outgrowth (Fig. 5).

Shortly after its birth, VD2 sends an anterior-projecting process along the ventral nerve cord. This process forms nascent presynaptic terminals to the ventral muscles as it extends (Fig. 5, Ventral). These nascent presynaptic terminals consist of swellings that contain presynaptic dense projections but either with no vesicles or with a few vesicles away from these projections (Fig. 5A-B). They first appear near the NMJs from the DD1 axon. The termini that are more proximal to VD2 soma appear smaller and less mature, suggesting that they continuously develop along the migrating process. As the larva grows, these swellings acquire vesicle pools and grow in size, making morphologically mature NMJs to the ventral muscles (Fig. 5C-D).

The migrating VD neurite next turns dorsally and joins the dorsal nerve cord (Fig. 5, Dorsal). As it enters the dorsal cord, it intercepts the postsynaptic field of embryonic DA and DB eMNs, its synaptic partners in the adult configuration (Fig. 5B-C). Targeting of the migrating VD2 neurite to its post-synaptic location is likely facilitated by DD1, which it wraps during commissural migration and extension along the dorsal nerve cord. Other VD iMNs use other structures to climb dorsally, but also wrap DD neurites in the dorsal nerve cord (Supplemental data sheets 1, 2). Migrating AS eMNs neurites similarly wrap existing neurites during outgrowth in the dorsal cord (Supplemental data sheets 3; Fig. 5 – Supp. 3c).

In the newborn larva, DD neurites are post-synaptic to embryonic DA and DB eMNs in the dorsal nerve cord. As the larva grows, VD2’s dorsal process gradually replaces DD1 in this apposition (Fig. 5, Dorsal; Fig. 5 – Supp. 2i-V). Thus, when the DA and DB eMNs contract dorsal muscles, the VD iMNs take over the original role of the DD iMNs in relaxing ventral muscles during dorsal bends.

We found it remarkable that the axonal fate of VD2’s ventral process is already established before its dendrite has formed. It is also striking that its neurite is already in appropriate apposition to future partners and develops synaptic structures while it extends. We observed similar morphological events that anticipate future wiring in the developing neurites of all post-embryonic eMNs (Fig. 5 – Supp. 3).

These observations suggest that the wiring of post-embryonic motor neurons is at least partly intrinsically programmed. Physical interactions between post-embryonic and embryonic iMNs may facilitate wiring maturation but may not be essential (see Discussion).

### Expression of post-synaptic receptors corroborates the temporal and spatial pattern of wiring transition

Serial EM reconstruction often identifies synapses by morphological features within presynaptic terminals, because many synapses, such as mammalian inhibitory synapses lack striking morphological features at the postsynaptic sites^20^. Post-synaptic partners are often identified by their close physical proximity to presynaptic structures^7,9,21^. Our characterization of the motor circuit remodeling process is contingent on reliability of a proximity-based postsynaptic annotation.

To verify our conclusions, we examined the developmental changes of post-synaptic receptors at iMN NMJs. UNC-49 is the *C. elegans* ionotropic GABAa receptor^22^, forming a GABA-gated chloride channel in the muscle cells that inhibits action potential firing^23–25^. Using endogenously tagged UNC-49::tagRFP^26^, we evaluated the consistency between its localization with the change of embryonic iMN’s post-synaptic partners by longitudinal fluorescent imaging (Methods) (Fig. 6A).

**Figure 6.**
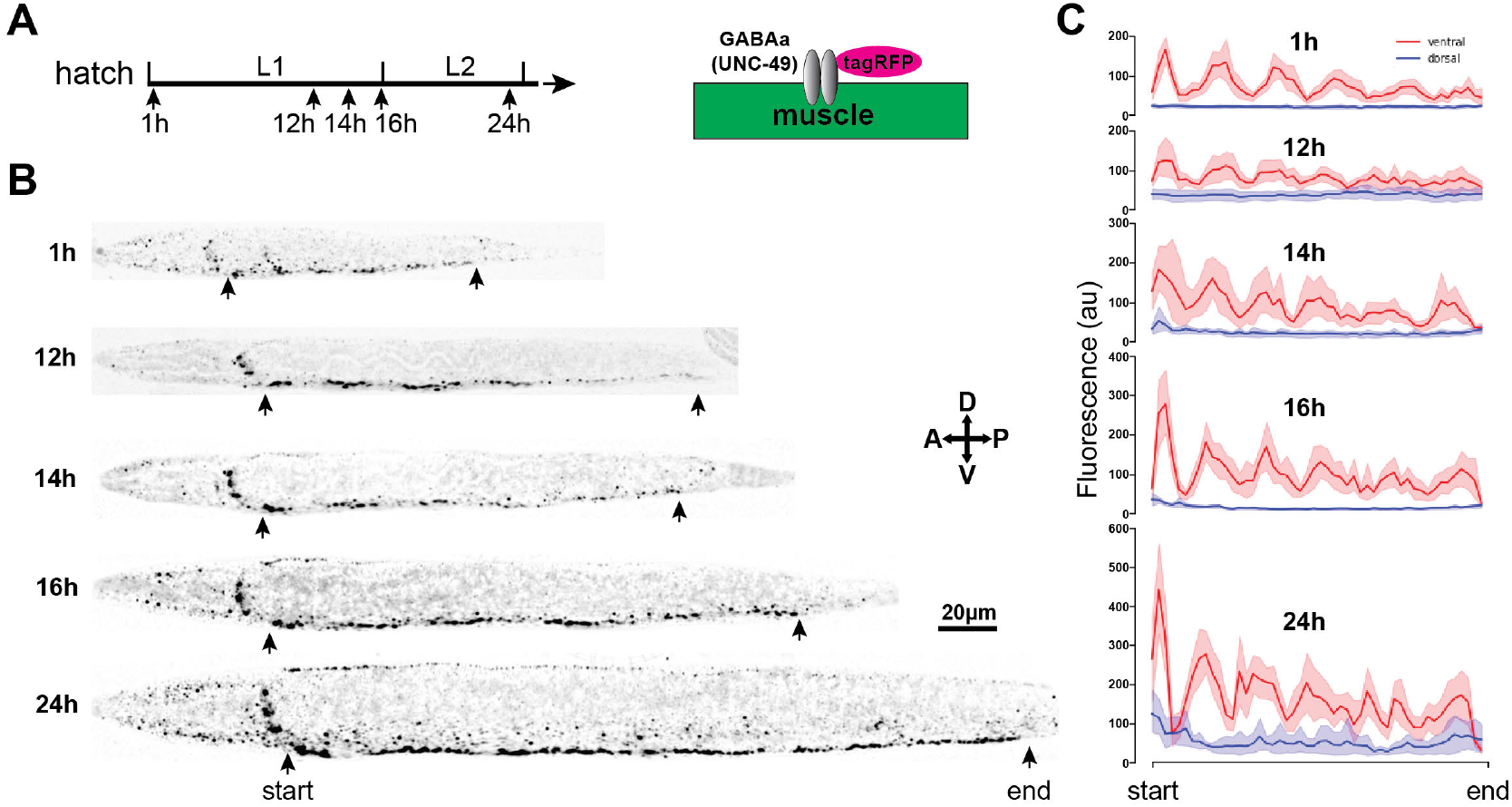
Receptor clustering at postsynaptic termini follows morphological synapse rewiring of iMN motor neurons. (A) Schematics of endogenously tagging of the postsynaptic GABAa receptors. (B) Example images (B) and quantitative representation (C) of the expression pattern of endogenous postsynaptic GABAa::RFP clusters in *C. elegans* larvae at different post-embryonic developmental ages. At early to mid-L1 stages, these clusters are presented along the ventral cord. These clusters remain prominent and continue to grow along the ventral cord while dorsal signals emerge gradually towards the late L2 stage.

At birth, iMNs make NMJs strictly to ventral muscle cells. Consistently, from early L1 to mid-L1 stage, endogenous GABAa receptors were present only along the ventral cord (Fig. 6B, C). They appeared as six distinct clusters, in correspondence to post-synaptic fields of clustered NMJs at the anterior axon of 6 DD neurons in our EM reconstruction (Fig. 4A).

At the late L1 stage, GABAa receptor clusters remained prominent along the ventral nerve cord, and weak signals appeared in the dorsal nerve cord, sporadically among and within individual larva (Fig. 6B, C). By the L2 stage, prominent ventral GABAa receptor clusters remained and became more evenly distributed throughout the cord, whereas dorsal signals also increased in intensity and spread along the dorsal nerve cord (Fig. 6B, C).

Emergence of dorsal clusters while maintaining prominent ventral clusters is consistent with characteristics of remodeling iMN neurons revealed by the EM reconstruction. When post-embryonic iMNs begin to build NMJs along the ventral nerve cord, embryonic iMNs maintain NMJs to ventral muscles so that they are never devoid of inhibitory inputs. Concurrently, embryonic iMNs begin to build NMJs to dorsal muscles.

## Discussion

Throughout life, a nervous system continues to change. This may occur as part of an animal’s developmental program, or as part of the adaptive mechanism to new experiences. These changes take diverse forms. Vertebrate brains, for example, typically maintain a stable cell population but substantially expand neurites and refine wiring after birth. The developing spinal cord, on the other hand, may require substantial addition of new neurons.

After birth, the *C. elegans* motor circuit undergoes significant changes in neuron number, neurite apposition, and synaptic connectivity. However, it maintains continuous locomotory activity and similar undulatory bending patterns^15,16,27^ (Fig. 1C, D). Here, we describe strategies that allow the motor circuit to dramatically change its structure and wiring while preserving its ability to drive locomotion (Fig. 7).

**Figure 7.**
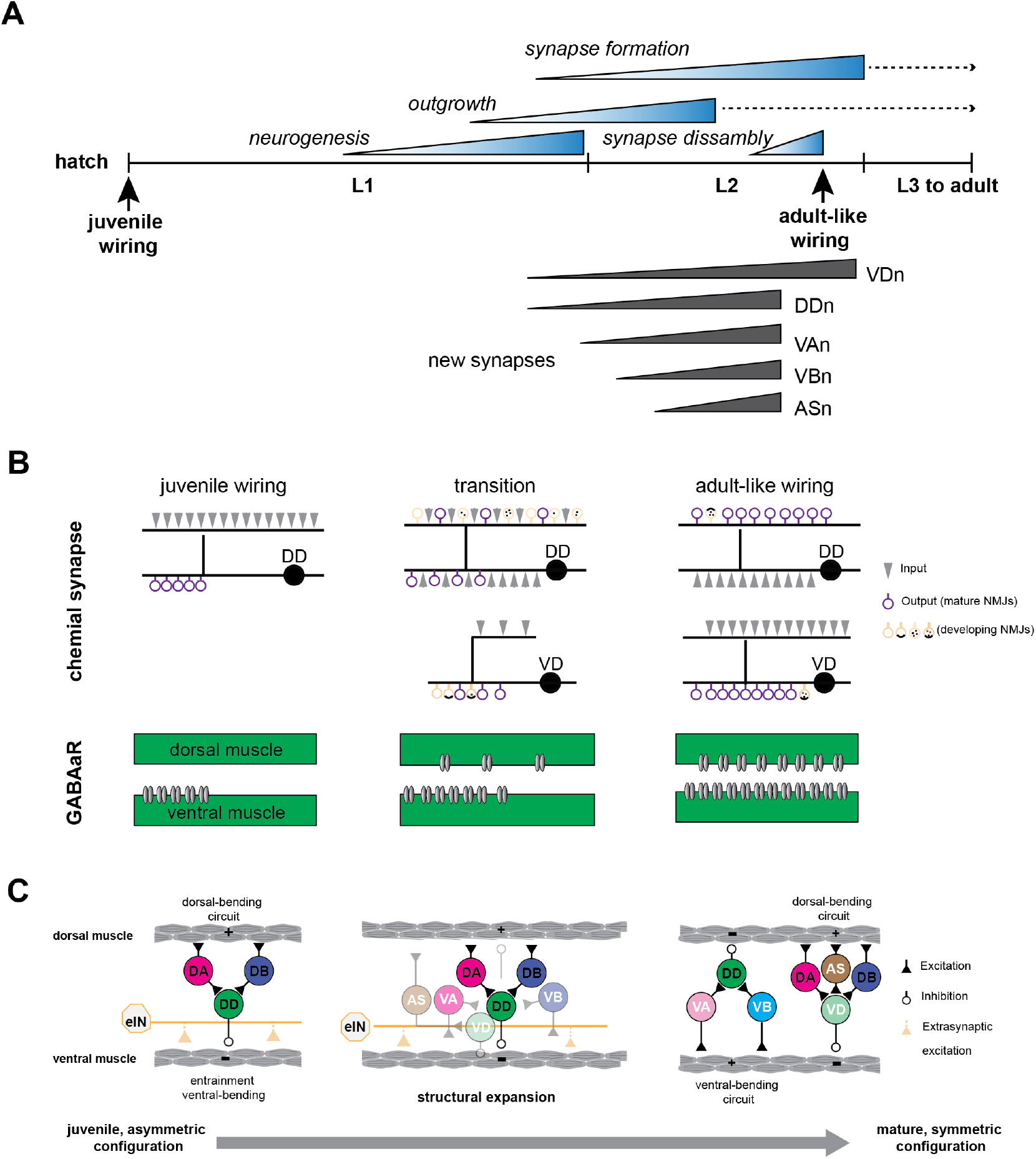
A summary of post-embryonic motor circuit maturation. (A) Staggered developmental timelines for anatomic events of motor circuit maturation. Above the timeline denotes the periods of post-embryonic motor neuron neurogenesis, neurite outgrowth, formation of new synapses, and disassembly of juvenile synapses. Below the timeline denotes the period of transition to adult wiring configuration by the neuron class. Adult-like wiring denotes a state where immature synapse structures are still be observed. Dashed lines indicate continued neurite elongation and synapse addition as the larva grows in body size. (B)*(Upper panels)* Rewiring of embryonic inhibitory motor neurons (DD) involve a switch of their axon and dendrite, during which both processes built mixed pre- and post-synaptic structures. Concurrently, post-embryonic VD iMNs take up DD’s juvenile wiring pattern as their neurites make NMJs as they extend. DD iMNs retain and grow their juvenile ventral NMJs, but begin to make new dorsal NMJs and to receive input from newly born post-embryonic eMNs, in anticipation for switching to the mature motor circuit configuration. Both build new NMJs sequentially but in opposite orders: DD accumulates the vesicle first; VD adds the presynaptic dense projection (active zone) first. New connectivity has been built before juvenile connectivity is disassembled. (*lower panels*) Post-synaptic receptor clusters (GABAa/UNC-49) at the iMN NMJs are present at ventral muscles in L1, in apposition to the anterior region of DD axons. As DD remodels to make NMJs to dorsal muscles, GABAa receptors start to appear in dorsal muscles while ventral clusters grow. In the mature configuration, GABAa clusters are presented in both ventral and dorsal muscles. (C)*(left)* The *C. elegans* motor circuit acquires structural and functional symmetry through post-embryonic development. Motor neurons of juvenile animals are asymmetrically wired to generate dorsal-bending. Juvenile animals achieve symmetric dorsal-ventral bending through extrasynaptic excitation from excitatory interneurons (eINs) to ventral muscle cells and entrainment by the dorsal bending circuit. *(Center)* During post-embryonic maturation, embryonic eMNs and iMNs form new wiring with post-embryonically born eMNs (VA, VB, AS) and new iMNs (VD) (grey arrows and circles with white labels). Embryonic iMNs (DD) may physically guide their replacement, the VD iMNs into the post-embryonic field of their juvenile partners. New circuitry is built while the juvenile connections are retained and functional (dark arrows). *(Right)* After the juvenile connectivity is disassembled, the mature motor circuit operates as two symmetrically wired sub-circuits, one for dorsal bending and one for ventral bending.

### Maturation generates symmetry

*C. elegans* is born with an asymmetrically wired motor circuit. During post-embryonic development, the motor circuit gains structural symmetry (Fig. 7).

Motor neurons of the newborn L1 larva are wired to contract dorsal muscles through excitatory motor neurons and simultaneously relax ventral muscles through inhibitory motor neurons, thus promoting dorsal bending. Ventral bending occurs through entrainment to the activity of the dorsal bending circuit^6,16^. In contrast, the adult motor circuit is symmetric, with separate circuits for dorsal and ventral bending^9^.

### An orchestrated and gradual transition

Post-embryonic motor neuron birth begins in the second half of the L1 stage and completes by the beginning of the L2 stage^5^. Neurite extension and wiring maturation take place gradually and continue throughout the larval stages. The conversion from the L1 to adult configuration occurs continuously along the body, in an anterior-to-posterior sequence that recapitulates the birth order of post-embryonic motor neurons (Fig. 7A).

After the birth of the post-embryonic motor neurons, the juvenile dorsal bending circuit remodels its wiring. The inhibitory motor neurons get recruited to a new ventral bending circuit, receiving input from newly born post-embryonic ventral excitatory motor neurons, and forming new inhibitory output connections with dorsal muscles - a reversal of their juvenile polarity. Meanwhile, post-embryonic inhibitory motor neurons take over the vacated inhibitory role in the dorsal bending circuit, receiving input from the embryonic dorsal excitatory motor neurons, and forming new inhibitory output connections with ventral muscles. These changes result in the generation of a new ventral bending circuit, and a substantially remodeled dorsal bending circuit (Fig. 7B, C).

Rewiring of the embryonic inhibitory motor neurons disconnects them from the dorsal bending circuit and connects them with the ventral bending circuit (Fig. 7C). This transformation takes the form of converting the ventral process from an axon to a dendrite, and the dorsal process from a dendrite to an axon (Fig. 7B). This conversion is gradual. First the dorsal process builds nascent pre-synaptic structures, while retaining its juvenile dendritic input. The ventral process begins to acquire input from newly born post-embryonic ventral excitatory motor neurons, while retaining its juvenile neuromuscular junctions to ventral muscle cells. Embryonic motor neurons only dismantle their juvenile connectivity when the adult-stage circuitry is morphologically mature.

Functional replacement of the juvenile dorsal bending circuit by the new dorsal and ventral circuits must be tightly orchestrated. One potential mechanism is intrinsic, preparatory rewiring. Embryonic inhibitory neurons build structures for the adult circuit while continuing to serve their role in the juvenile circuit. Another potential mechanism is extrinsic, communicative rewiring. The embryonic inhibitory motor neurons serve as guideposts to place the neurites of post-embryonic inhibitory motor neurons that will later take over their circuit function. Points of physical contact between the embryonic and post-embryonic inhibitory motor neurons are where the embryonic excitatory motor neurons begin to exchange post-synaptic partners.

These mechanisms allow a seamless transition from the newborn to the adult-stage circuit with a structural and functional symmetry (Fig. 7C).

### Consistency with previous studies

Our observations of post-embryonic motor circuit maturation confirm and extend insights from previous electron and light microscopy studies.

The initial discovery of the DD embryonic motor neuron’s neurite and wiring remodeling was made through a partial EM reconstruction of the mid-L1 larva when post-embryonic neurogenesis had just begun^6^, and validated by the position of fluorescent markers that label the DD NMJs^8,28,29^. By following a presynaptic marker, remodeling of their synaptic output was found to occur in an anterior-to-posterior order, completing by the L2 or L3 larva stages^8,11^.

Our new EM datasets, with reconstructions of the entire motor circuit at developmental ages before, during and after post-embryonic neurogenesis consolidate these insights. Our datasets further offer a full account of the temporal and spatial ordering of maturation events of all motor neurons from neurite outgrowth to synaptic wiring. Owing to a better sample preservation method^21,30,31^, these datasets reveal sequential ultra-structural events during the assembly of wiring across this developing circuit. These observations establish a foundation for mechanistic dissection of each step of the circuit remodeling.

### Reversing axon-dendrite polarity without pruning

Embryonic inhibitory motor neurons fully reverse their neurite’s axon-dendrite identity without breaking them down. This implies an intrinsic ability to compartmentalize and simultaneously accommodate the axon and dendritic properties in the same neurite. Unipolar neurons that intercalate axonal and dendritic regions in the same neurite exist in many species from *C. elegans* to mammals. The assignment of axonal and dendritic compartments might be achieved by manipulation of intracellular transport.

Consistent with this idea, the embryonic inhibitory motor neurons require multiple kinases that play a general role in cell polarity to restrict the presynaptic machinery to the axonal process^32–35^. Blocking microtubule dynamics^36^ and removing an ER-nuclei protein^37^ also block the reversal of DD axon-dendrite polarity.

### Intrinsic programming with flexibility

Temporal and spatial orchestration of post-embryonic motor circuit maturation implies that many events are intrinsically programmed. Migrating neurites of post-embryonic neurons initiate synapse development with their future partners as they extend. Thus, the axon-dendrite identity as well as wiring partners are likely encoded by intrinsic developmental programs.

Following observations also suggest that both embryonic and post-embryonic motor neurons are programmed for wiring and re-wiring.

In the *unc-55* lineage mutant where post-embryonic inhibitory motor neurons do not properly differentiate^38^, embryonic inhibitory motor neurons still innervate dorsal muscles in adults. In another lineage mutant *lin-6/mcm-4*, precursors for post-embryonic motor neurons fail to divide^39^, and all post-embryonic motor neurons are absent^6^. In a late stage (L4) larva, embryonic motor neurons remodel NMJs to dorsal muscles without presynaptic input to their ventral processes^6^. Consistently, their presynaptic marker eventually switches from the ventral to dorsal processes^8^. Lastly, when all embryonic inhibitory motor neurons are ablated, post-embryonic motor neurons still innervate ventral muscles in adults^38,40^. Thus both embryonic motor neurons and post-embryonic motor neurons can wire independently.

A transcription factor UNC-30 intrinsically specifies the fate of GABAergic neurons^41^. In an adult *unc-30* mutant, inhibitory motor neurons exhibit aberrant soma position, neurite trajectories, and synaptic wiring, with input most severely disrupted (Extended Figure 1 in^13^). Further, the final wiring pattern of either embryonic or post-embryonic inhibitory motor neurons may not require synaptic transmission, because the switch of presynaptic and postsynaptic markers for embryonic motor neuron’s NMJs occurs normally in the absence of GABAergic signaling^36^.

Some aspects of embryonic motor neuron’s remodeling may require post-embryonic motor neurons.

In the L4 stage *lin-6/mcm-4* mutants, their dorsal process appears partially remodeled: making NMJs to dorsal muscles and maintaining some postsynaptic input from the L1 circuit, presumably due to the absence of anticipated new post-synaptic partners^6^. Neurexin/NRX-1, an adhesive molecule is required to induce postsynaptic receptor clustering in the remodeling dendrite^42^. NRX-1 is specifically required in cholinergic motor neurons^42^, raising the possibility that their adult input partners, excitatory post-embryonic motor neurons, are involved in this process.

Intrinsic programming also does not preclude a circuit-level flexibility during post-embryonic maturation. By visualizing embryonic inhibitory motor neurons with fluorescent synaptic markers, many genes have been found to alter the timing of their rewiring. Two lineage mutants *lin-14* and *lin-4* accelerate and delay the timing of global post-embryonic development events, including the rewiring^8^. Without post-embryonic neurons, embryonic motor neurons significantly delay the switch of their presynaptic input from ventral to dorsal muscles^8^, and may fail to fully remove the L1 stage postsynaptic input^6^. Thus, while neither neuron class requires each other to form NMJs with their muscle targets in the adult stage, physical interactions between embryonic and post-embryonic motor neurons may promote their remodeling. Timing of their rewiring might also be slightly delayed with reduced synaptic transmission or neuronal activity, or accelerated with increased vesicle release^43–45^.

We propose that physical communication and neuronal activity are facilitatory. They fine-tune an intrinsic developmental program to achieve coordination, flexibility and adaptability.

### Robustness of the motor circuit

A gradual and orchestrated remodeling of the L1 motor circuit to its adult form explains a seamless transition without functional disruption. Understanding how the circuit maintains the same motor pattern leads to another question. How does the wiring of the L1 motor circuit, which resembles half of the adult motor circuit specific for dorsal bending, generate both dorsal and ventral bending?

In a separate study, we resolved this question: before post-embryonic neurons are born, extrasynaptic cholinergic signaling from excitatory premotor interneurons and extrasynaptic GABAergic signaling from inhibitory motor neurons functionally compensate for the lack of a structural ventral bending circuit^16^. Thus, an adult-like motor pattern is already in place before the adult motor circuit develops.

With mechanisms that functionally compensate for the newborn’s asymmetric structure in place, a gradual replacement of adult-stage wiring can occur along the anterior-posterior axis without causing changes to motor patterns. This explains how genetic mutants that alter the timing of remodeling can nevertheless maintain an undulatory motor pattern.

These mechanisms work in concert to lend robustness to the developing motor circuit, ensuring continuity of motor function during structural changes. Robustness benefits survival in presence of genetic and environmental adversity.

### Why does it remodel

If motor patterns do not change from the L1 to adult, why remodel at all? This might be a strategic decision by *C. elegans*. Driven by its short life cycle, it needs to optimize the relative amount of circuit maturation that occurs in the embryonic and post-embryonic stages.

With rare exceptions, post-embryonic neurogenesis in the *C. elegans* is dedicated to either expanding the motor circuit during early larva stages or developing sex-specific circuitry in later larva stages. The structural symmetry of the motor circuit is acquired through post-embryonic neurogenesis and wiring remodeling. By contrast, at birth, the brain has completed neurogenesis, neurite formation, and neuropil assembly. An embryo may prioritize the development of its brain over its motor circuit so the animal is able to make adaptive decisions, essential for survival at birth.

But motility is also critical for survival. The animal likely develops robustness with compensatory mechanisms for an unfinished motor circuit, such that its structural changes make little impact on functions. Having two different functional mechanisms to generate the same motor pattern, a gradual structural transition between their respective configurations enables a transition without behavioral disruption.

### Different approaches by the brain and body

All neurons share the ability to modulate synapses, changing their physical size, transmission strength, and molecular composition for signaling in response to development and stimulation^46^. Synapse plasticity has long been viewed to underlie learning^47^. However, adaptive and changing behaviors reflect plasticity at the systems level^48^. Synaptic plasticity should be orchestrated across circuits and development.

One needs to ask how intact circuits coordinate their structural and functional changes in response to development and experience. This allows meaningful comparison of synaptic plasticity across circuits and species. For *C. elegans*, the brain and body circuits take different approaches to reach maturity. The anatomy of the brain is already mature at birth. Wiring simply expands on a maintained topology of neuropil and may support a maturing and adaptive behavioral repertoire^21^. The body on the other hand strives to maintain the same output, while its anatomy and wiring dramatically expands and remodels. Their structural-functional relationships serve different ecological needs. These needs drive diverse approaches for plasticity or robustness in underlying circuits.

## Star Methods

### RESOURCE AVAILABILITY

#### Lead contact

Further information and requests should be directed to lead contacts.

#### Data and code availability

Original electron micrograph data will be freely available at bossdb.org. MATLAB scripts for calcium imaging analyses are available in GitHub https://github.com/zhen-lab/Beta\_Function\_Analysis. Original light microscopy data will be made available upon request to lead contacts.

### EXPERIMENTAL MODEL AND SUBJECT DETAILS

*C. elegans* strains were grown and maintained on nematode growth media (NGM) plates seeded with the *Escherichia coli* strain OP50 at 22.5 C. The N2 Bristol strain was obtained from *Caenorhaditis* Genetics Center (CGC). The EN296 [*UNC-49::TagRFP*] strain^26^ was obtained from the Bessereau lab. The AQ2953 (*ljIs131[Pmyo-3-GCaMP3::UrSL2::RFP]* strain^49^ was obtained from the Schafer lab.

### METHOD DETAILS

#### The age of a larva

Because the rate of developmental changes is dependent on raising temperatures, the age of larva (hours after hatching) is scaled to the lineage chart (N2 animals raised at 20°C) using established post-embrynic anatomical landmarks^5^. For electron microscopy studies, see Table 1 for main landmarks used to determine the age of larva. For the age of non-wild-type animals (muscle GCaMP::RFP and UNC-49::RFP), we scaled to the lineage chart by two landmarks: the presence and absence of L1 alae, and the number of gonad cells revealed by DIC imaging.

#### Muscle calcium imaging in crawling larva

Animals expressing GCaMP3 and RFP in body wall muscles (*ljIs131[Pmyo-3-GCaMP3::UrSL2::RFP]*^49^) were synchronized, sandwiched between a 2% agarose pad and a glass coverslip. Animals were imaged using a compound fluorescence microscope equipped with a dualbeam DV2 beam splitter to simultaneously record fluorescence from green and red channels. A motorized stage and custom plugin for μManager were used to track animals as they moved around the agarose pad, with a 10x objective and 10Hz sampling rate^50,51^. Body curvature and fluorescence intensity along the body axis were computed using custom scripts (^16^). Briefly head and tail are manually assigned in the first frame of the recording and tracked afterwards. The contour of the worm is divided into 100 segments along the anterior-posterior axis and further segment into the dorsal and ventral segments across the mid-line. Averaged intensity for pixels within each segment is used as proxy for the activity for this muscle segment. The phase difference was calculated by first using the Hilbert transform as implemented by MATLAB’s hilbert function to obtain an analytical representation of the signal. The difference between the phase angles obtained from the analytical representation of muscle activities and curvatures was used to generate phase difference histograms.

#### Swimming

For swimming assays, animals were placed in a drop of M9 on a glass coverslip, contained within a 1 cm diameter vaseline stamp, and imaged at 10Hz on a Zeiss V16 dissecting microscope. Swim cycles were manually counted from the videos.

#### Serial-section electron microscopy

Animals were prepared by high pressure freezing followed by freeze substitution in acetone containing fixatives^30,31^. The freeze substitution protocol was: -90°C for 96h in 0.1% tannic acid and 0.5% glutaraldehyde; wash with acetone 4x over 4h; exchange with 2% OsO4 and ramp to -20°C over 14h; hold at -20°C for 14h; ramp to 4°C over 4h; wash with acetone 4x over 1h. Freeze substituted samples were infiltrated with Spurr-Quetol resin, embedded as single animals and cured in a 60°C oven for 24h.

The 1-5hr and 15h animals were processed for TEM: serial 50nm (1-5hr) and 70nm (15hr) sections were collected on 2×0.5mm slot grids, postained with 2% UA and 0.1% lead citrate, and imaged at 0.7-1nm/pixel resolution using a Technai T20 TEM with AMT 16000 and Gatan Orius cameras. The rest of the animals were processed for SEM: serial 30nm sections were cut collected onto kapton tape using an ATUM^52^, glued to silicon wafers, post-stained with 4% UA and Leica Ultrostain II (3% lead citrate), and semi-automatically imaged using a FEI Magellan scanning electron microscope at 1-2nm/pixel^53^.

Micrographs from TEM and SEM were stitched into 3D volumes using TrakEM2^54,55^ and exported to CATMAID^56^ for cell identification, skeleton tracing and connectivity mapping^21,31^. All cells were identified in these datasets, and all neurites were traced through dorsal, ventral, circumferential and sub-lateral nerve cords.

For all EM micrographs, identity of all membrane profiles were identified, but only annotation for cells of the motor circuit (Figure 1B) and neurons and non-neuronal cells of consistent adjacency with motor circuit neurons (e.g. muscle arms, AVL, etc.) were shown to simplify presentation. Annotation for all cells in all our EM datasets will be made publicly available.

Volumetric reconstructions were performed manually in VAST^57^ and the resulting 3D models were processed using Autodesk 3ds Max or Blender. The 16hr dataset was realigned for improved volumetric reconstruction using a custom stitching pipeline^21^.

### Fluorescent microscopy of UNC-49

The EN296 strain^26^, with endogenously tagged *unc-49::tagRFP*, was fixed on ice using 2% paraformaldehyde and 1x modified Ruvkun’s witches brew^58,59^ to reduce gut autofluorescence. Endogenous fluorescence was imaged on a Nikon confocal microscope using images stacks followed by maximum intensity projections. For quantification of UNC-49::tagRFP fluorescence in the ventral and dorsal nerve cords, ROIs were drawn along the nerve cords in Fiji^54^ and fluorescence intensity along the body axis was plotted using a custom script that binned the ROI into 50 segments and computed mean intensity of each segment.

## Acknowledgements

We thank the CGC, William Schafer and Jean-Louis Bessereau for strains, Albert Cardona for help with TrakEM2 and CATMAID, many undergraduate students for contribution to the EM reconstruction, and John White for comments on the manuscript. BM was supported by the Mount Sinai Foundation; JKM was supported by National Science Foundation Physics of Living Systems (NSF 1806818); ADC was supported by the National Institute of Health (R35 GM134970 and R01 NS093588); JWL was supported by the National Institute of Mental Health, Silvio Conte Center (P50 MH094271), the National Institutes of Health (U24 NS109102-01), and the Multidisciplinary University Research Initiative (GG0008784). JWL, ADTS, JLB and MZ were supported by the Human Frontier Science Program (RGP0051/2014). ADTS and MZ were supported by the National Institutes of Health (R01-NS082525-01A1). ADTS was supported by National Institutes of Health Brain Initiative (1U01NS111697-01) and National Science Foundation BRAIN EAGER (IOS-1452593). MZ was supported by Canadian Institutes of Health Research (MOP-123250 and Foundation Scheme 154274), the Radcliffe Institute for Advanced Studies, and the Mount Sinai Foundation.

## Author contributions

MZ conceptualized and supervised the study with BM, DW, ADTS and JWL; BM, DW, JM, RS, DH, ADC and MZ generated, analyzed and interpreted data; DB, YW, YL and TA developed tools for data analyses or visualization; BM and MZ wrote the paper; ADTS and JWL edited the paper; all authors read and approved the paper.

## Supplemental information

**Table 1.**
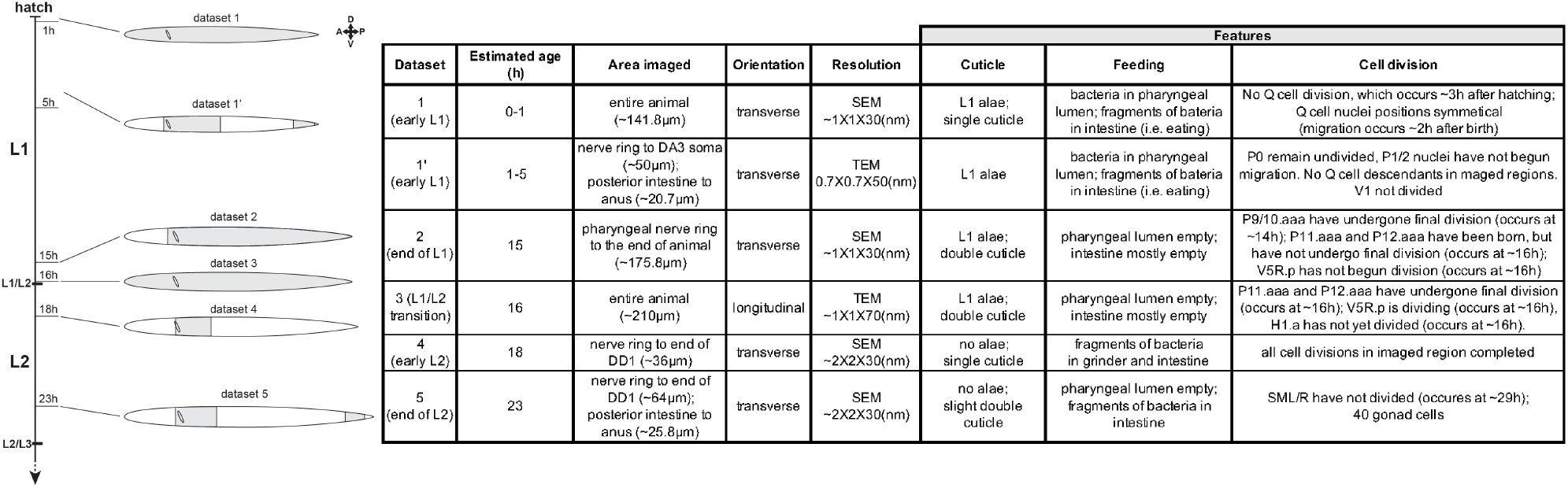
Anatomic landmarks for age-assignment of larva EM datasets.

**Figure 1 – Supp. 1.**
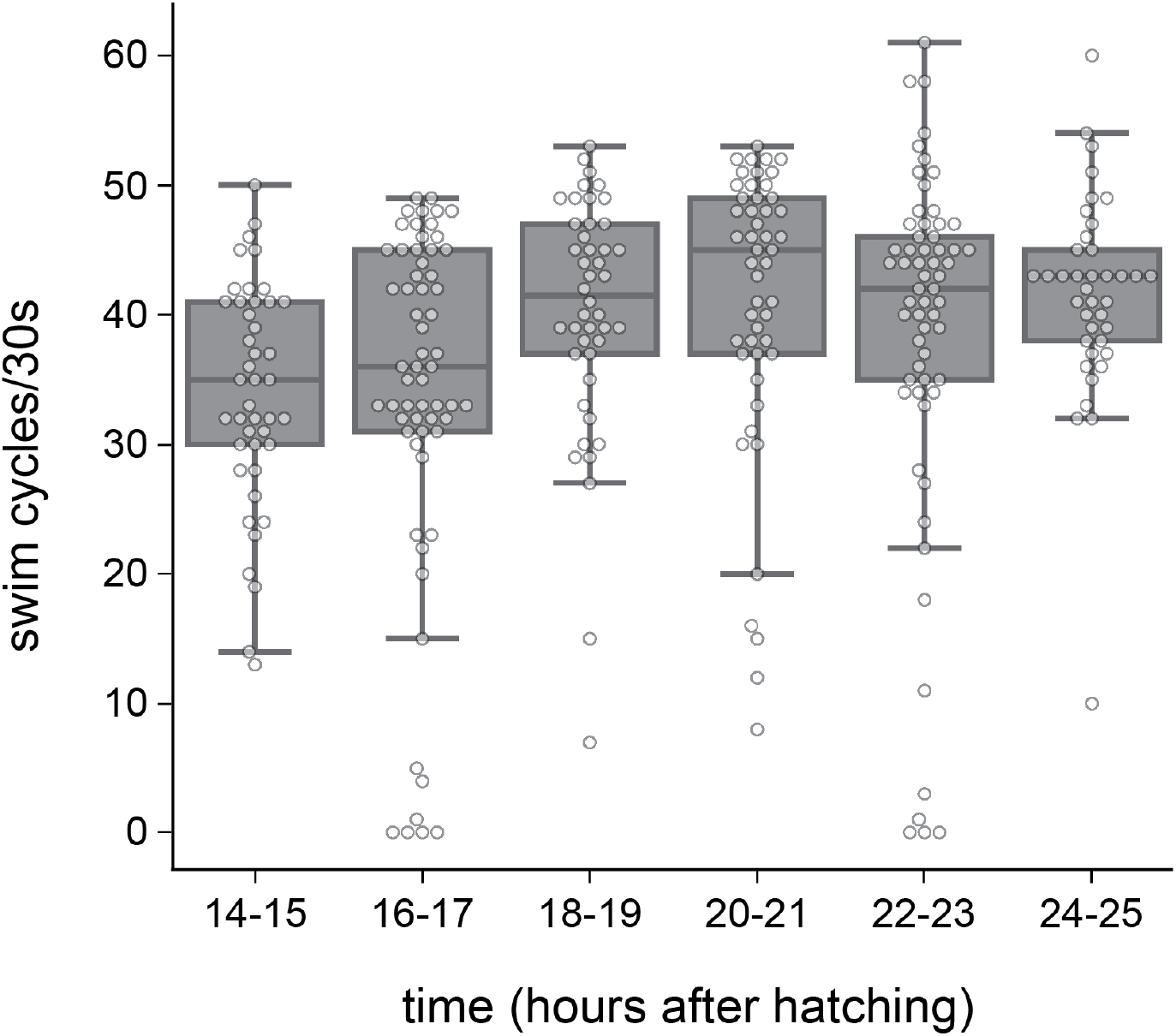
*C. elegans* larvae execute coordinated swimming before, during and after the juvenile-to-mature motor circuit transition.

**Figure 2 – Supp. 1.**
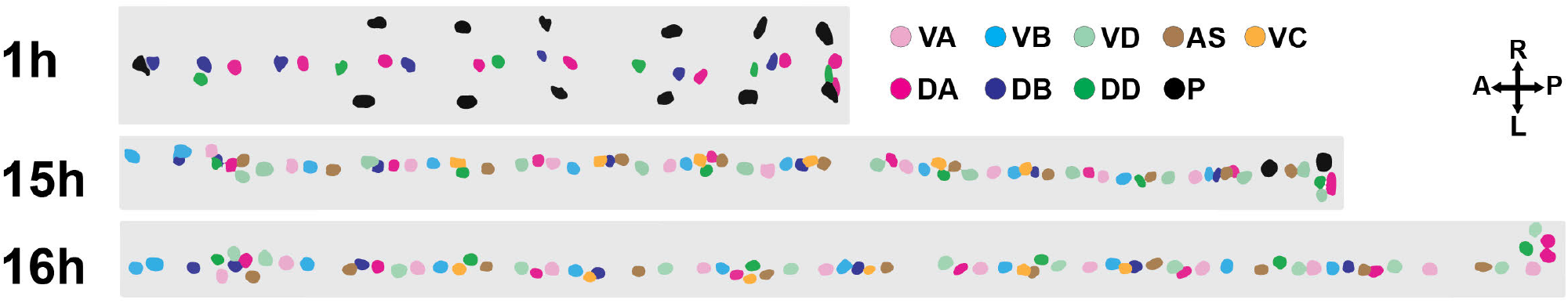
Body motor neurons and their precursors across the first larval stage. Nuclei of the body motor neurons and their precursors, including the VC class, in early L1 (1h), late L1 (15h) and the L1/L2 transition (16h).

**Figure 2 – Supp. 2.**
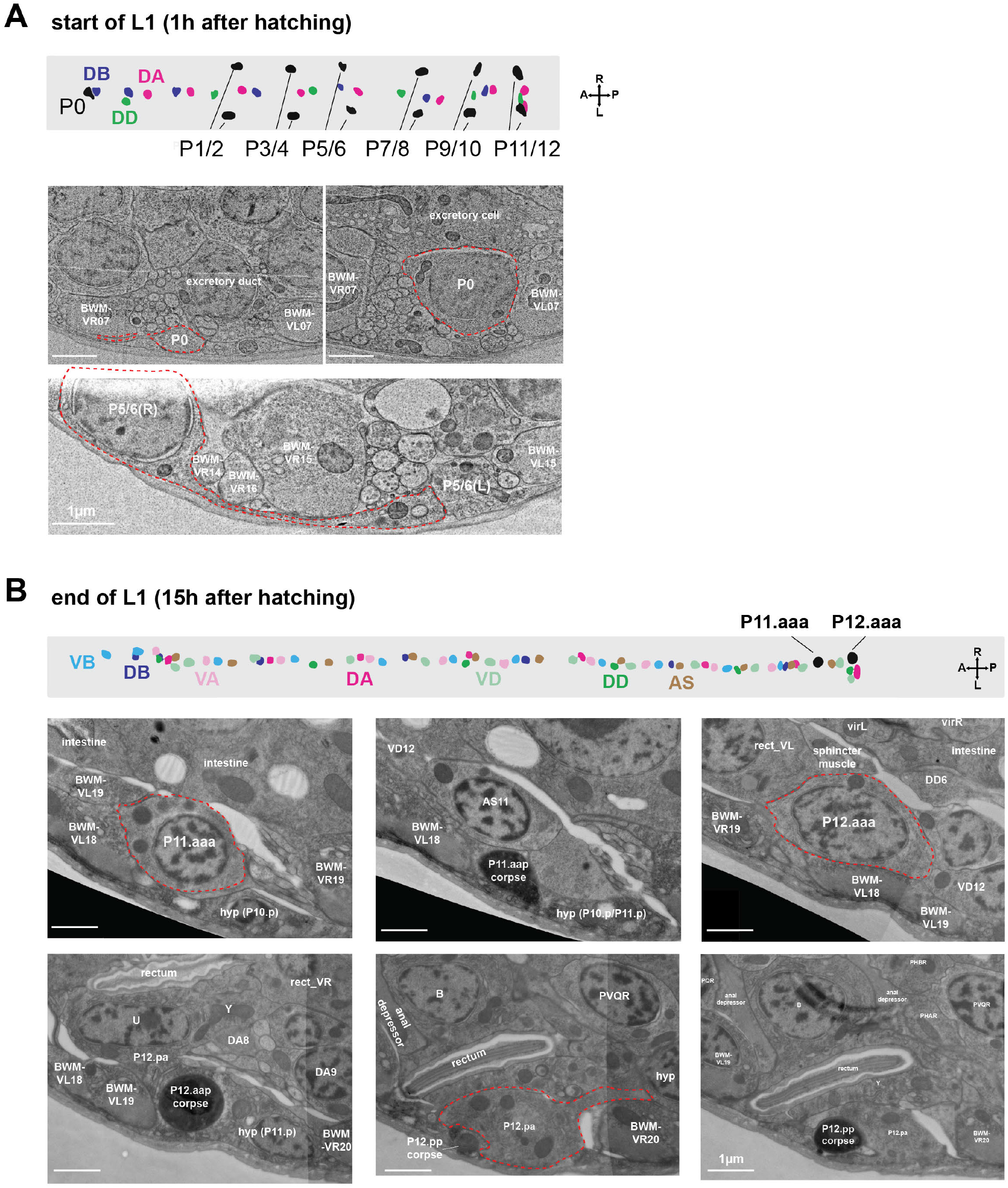
Morphology and migration of the precursor (P) cells that give rise of post-embryonic motor neurons. (A)*Upper panel* Nuclei of motor neurons in the ventral nerve cord of the newborn (L1) larva. The three embryonically born motor neurons are color-coded blue, green and magenta, respectively. P denotes the nuclei of precursor cells that will give rise post-embryonic neurons; *lower panel*, example electron micrographs of P cells and their processes. At birth, nuclei of P cells reside laterally and have not started migration, but their processes are already in the vicinity of the ventral nerve cord in their capacity as epidermal cells during early development (dashed red lines). (B)*Upper panel* Nuclei of motor neurons in the ventral nerve cord of the late L1 stage larva. All P cells have migrated to the ventral nerve cord, and most have completed multiple rounds of division that gives rise to post-embryonic motor neurons, color-coded with light blue, light magenta, light green, and brown, respectively. The only exception is P11 and P12, which have not finished their last round of division, leaving two neuronal precursors, P11.aaa and P12.aaa, in the ventral nerve cord. *Lower panel* Example electron micrographs of progeny of P11 and P12 cells, including those that are undergoing apoptosis. BWM: body wall muscle.

**Figure 2 – Supp. 3.**
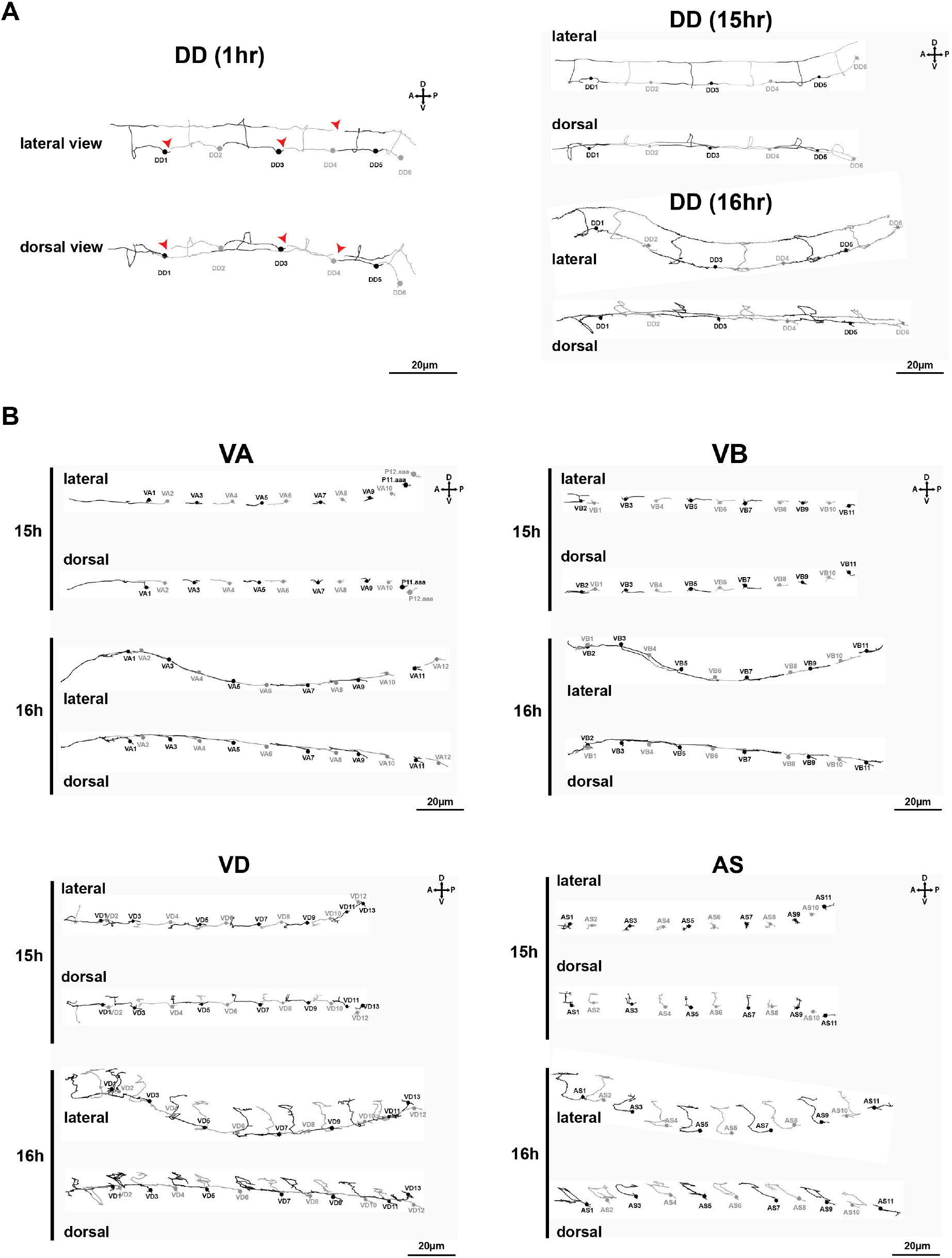
Skeleton reconstruction of all body motor neurons in early L1, late L1 and at the L1/L2 transition. (A) Left panel, neurites of embryonic motor neurons (DD1-DD6) do not fully tile the dorsal and ventral nerve cord at early L1 (1hr after hatching). Red arrows denote the physical gaps between neighbouring neurites. But each DD neuron makes at least one contact with its anterior neighbour and posterior neighbour through either the dorsal or ventral neurites. Right panels, by the end of L1 (15hr and 16hr), all dorsal and ventral neurites of DD1 to DD6 are connected with their neighbour, completing the tiling along the ventral and dorsal cord. (B) Neurites of the post-embryonic motor neurons (VA, VB, VD, AS) as they undergo outgrowth at the end of L1 stage (15hr) and the L1/L2 transition (16hr). Both dorsal and lateral views are shown.

**Figure 2 – Supp. 4.**
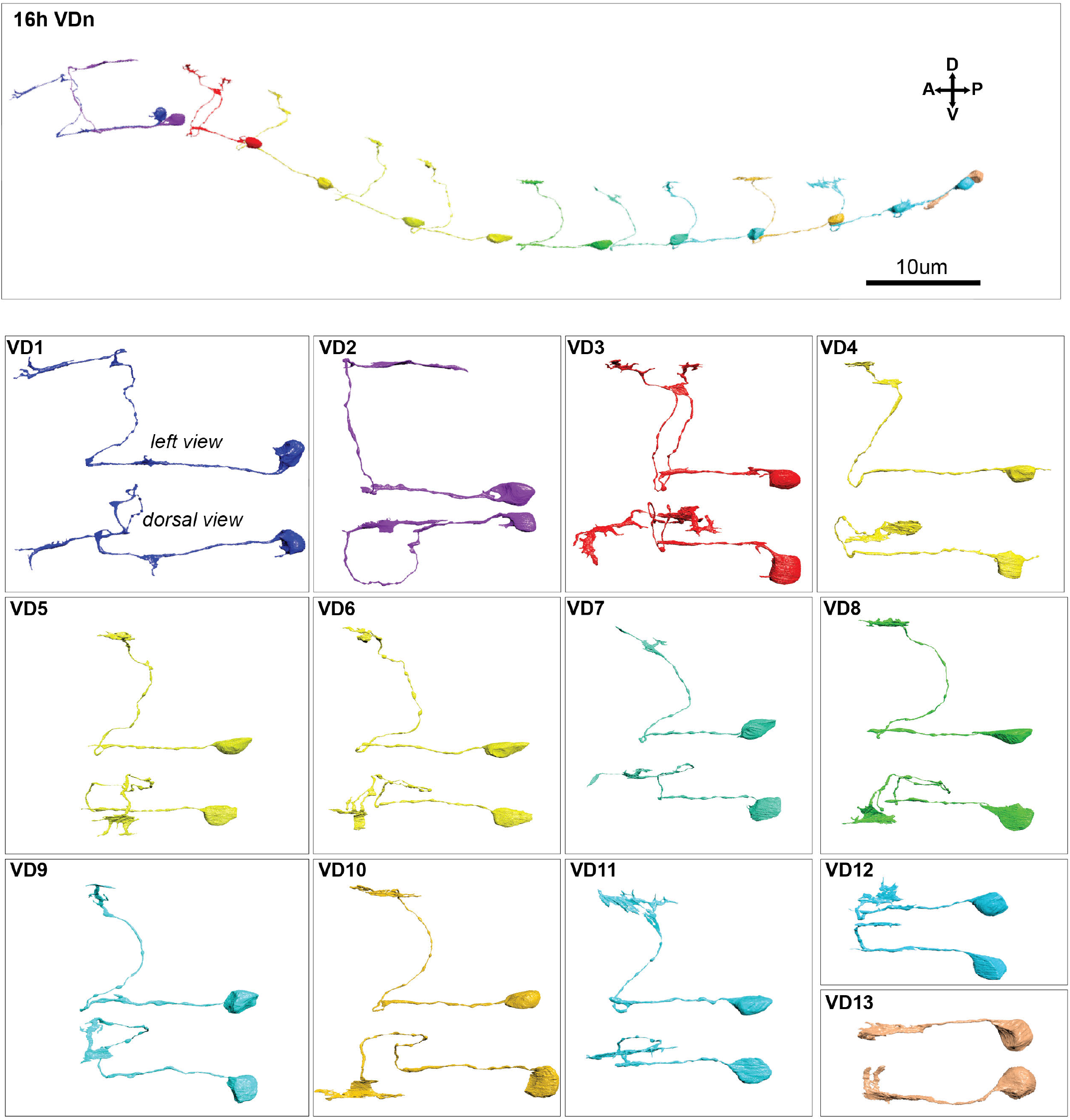
Volumetric reconstruction of the growing VD motor neurons at the L1/L2 transition. Neurons were reconstructed from serial micrographs, and are shown together (top) and separately (bottom) from lateral and dorsal perspectives. Outgrowth of the VD motor neurons follows roughly an anterior-to-posterior order, similar to their birth order, with anterior neurons at a more advanced stage of outgrowth.

**Figure 3 – Supp. 1.**
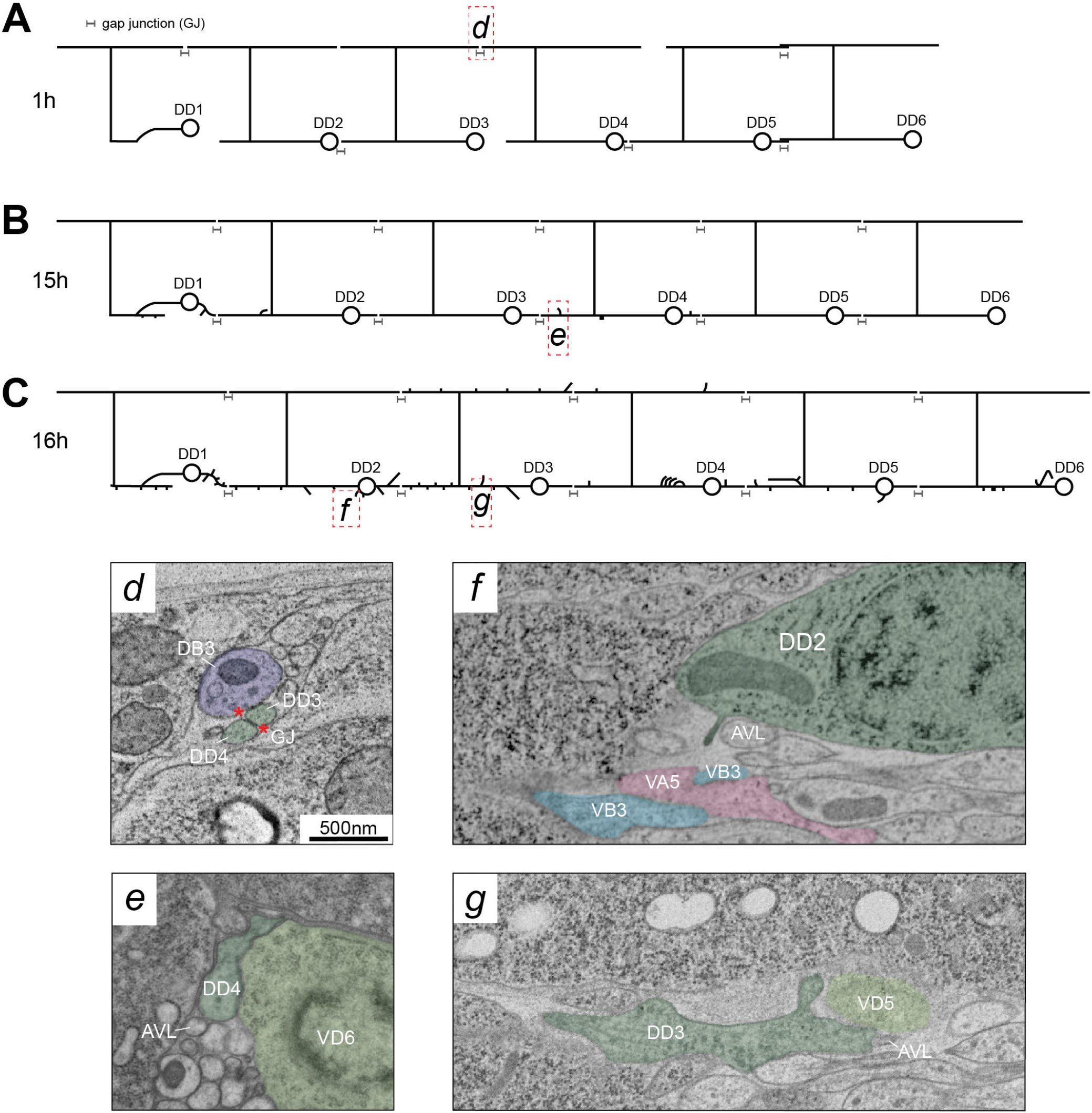
Examples of DD post-embryonic neurite outgrowth: tiling, guideposts and neighbourhood. Embryonic DD neurons continue their neurite outgrowth that serves two purposes. First is to complete tiling. (A) At birth, their neurites do not fully tile either dorsal or ventral cords, but each DD has at least one gap junction contact (d) with the neurite from its immediate anterior and posterior neighbours; GJ, gap junctions, bordered by red *. (B) By the end of L1 stage, DD neurites have fully tiled the dorsal and ventral cords, with gap junction connections between neurites of the neighbouring DD. Starting before the end of late L1 (B) and L1/L2 transition (C) stages, DD extends new branches that run adjacent to post-embryonic VD (e, g) and another GABAergic neuron AVL (e, f, g). Some ventral branches (f) are projecting dendrites towards the developing post-embryonic VA and VB eMN neurites.

**Figure 3 – Supp. 2.**
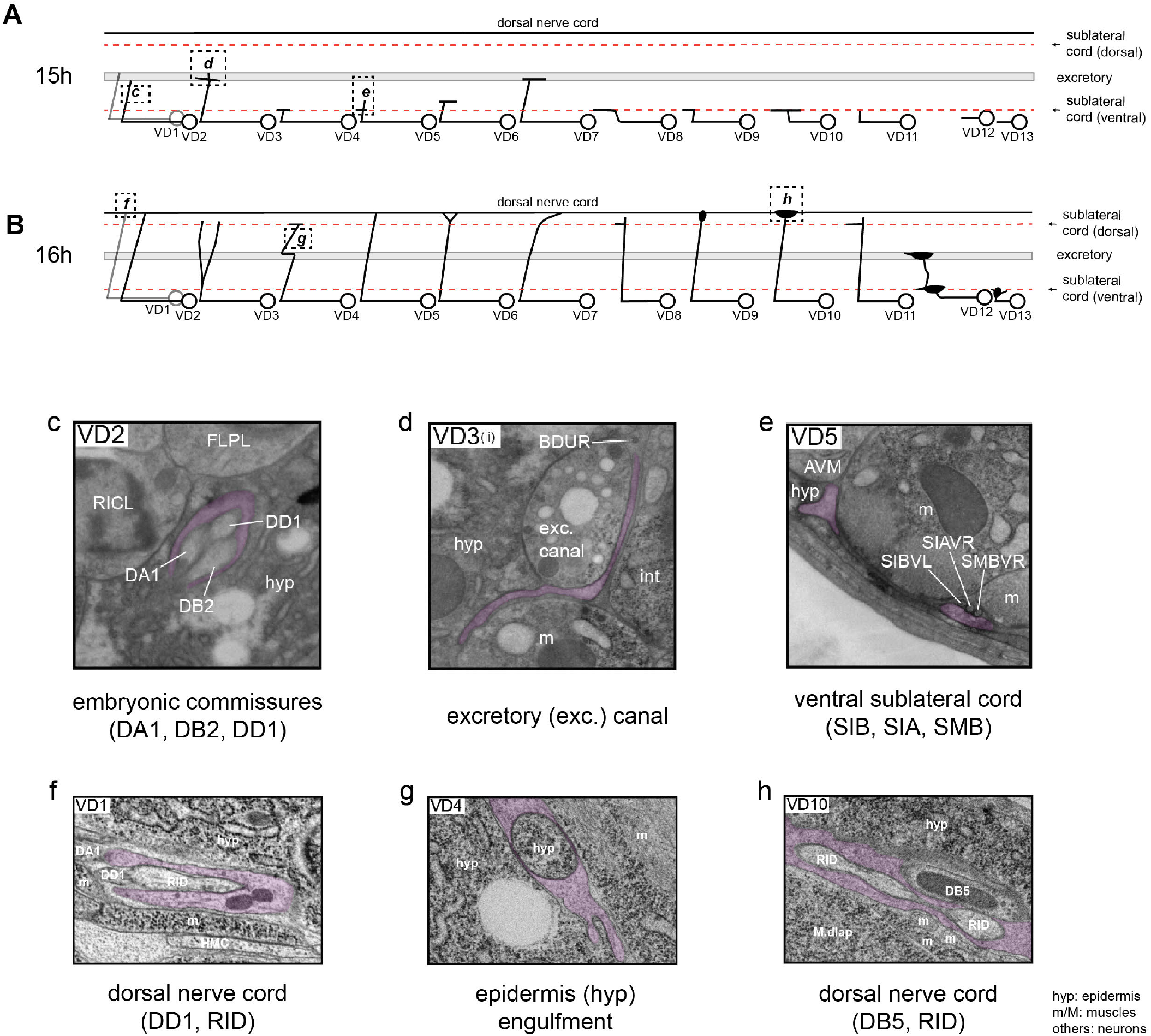
Examples of guideposts during VD post-embryonic neurite outgrowth. During outgrowth in the ventral and dorsal nerve cords, as well as during extension of the circumferential commissural processes, VD neurites interact with structures including the sublateral cords, the excretory canal, and DD motor neurons. Cartoons illustrating neurite morphology are shown for late L1 (15h, *top*) and the L1/L2 transition (16h, *bottom*), along with representative electron micrographs that illustrate some of the characteristic features exhibited during VDn outgrowth.

**Figure 4 – Supp. 1.**
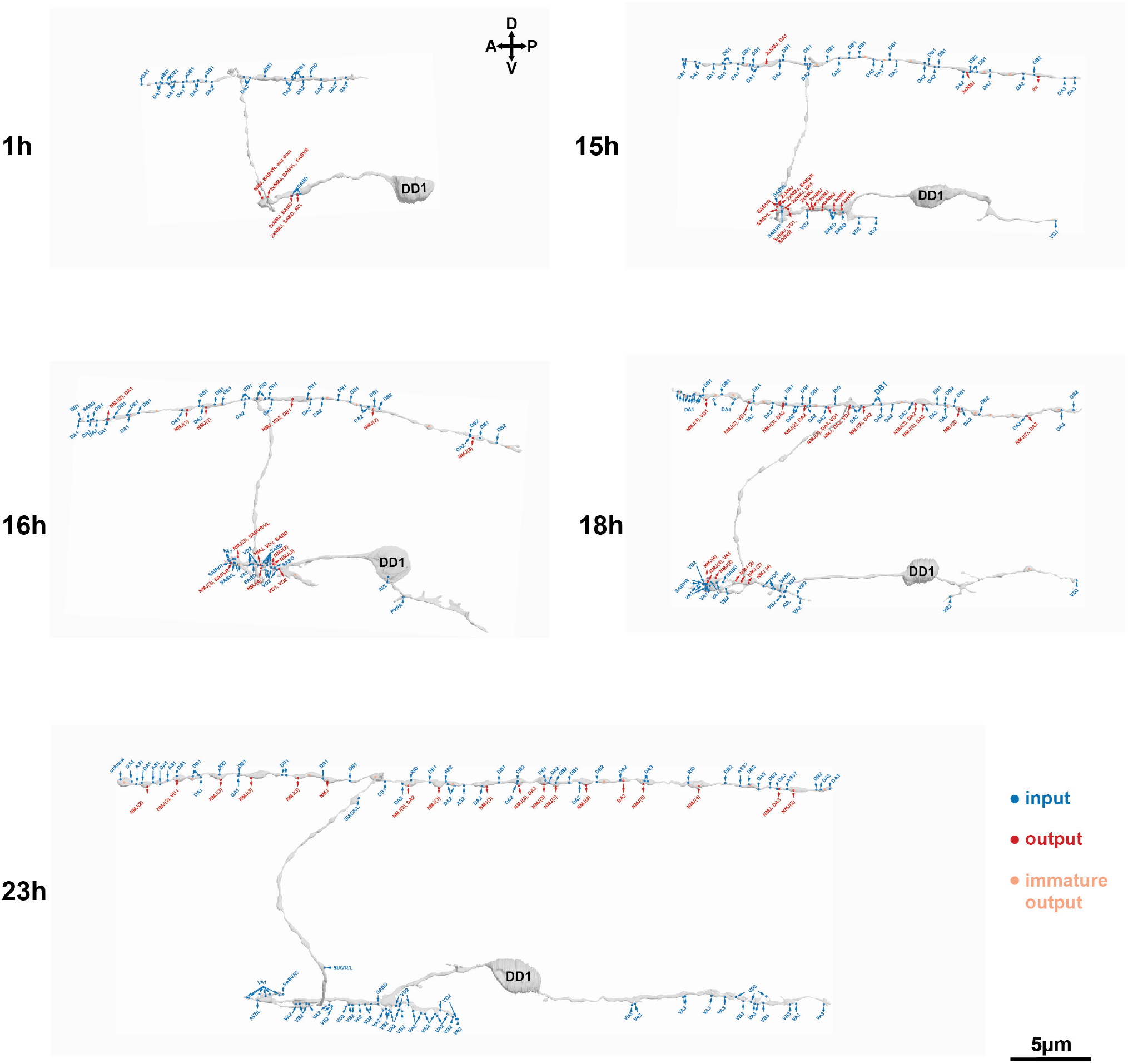
Connectivity of DD1 before, during and after remodeling. Developmental wiring diagram of DD1 at early L1 (hr), late L1 (15hr), L1/L2 transition (16hr), early L2 (18hr) and late L2 (23hr). NMJ, neuromuscular junctions. Numbers in brackets denote the number of chemical synapses for both input and output cells. All inputs and outputs without a number have a single chemical synapses.

**Figure 4 – Supp. 2.**
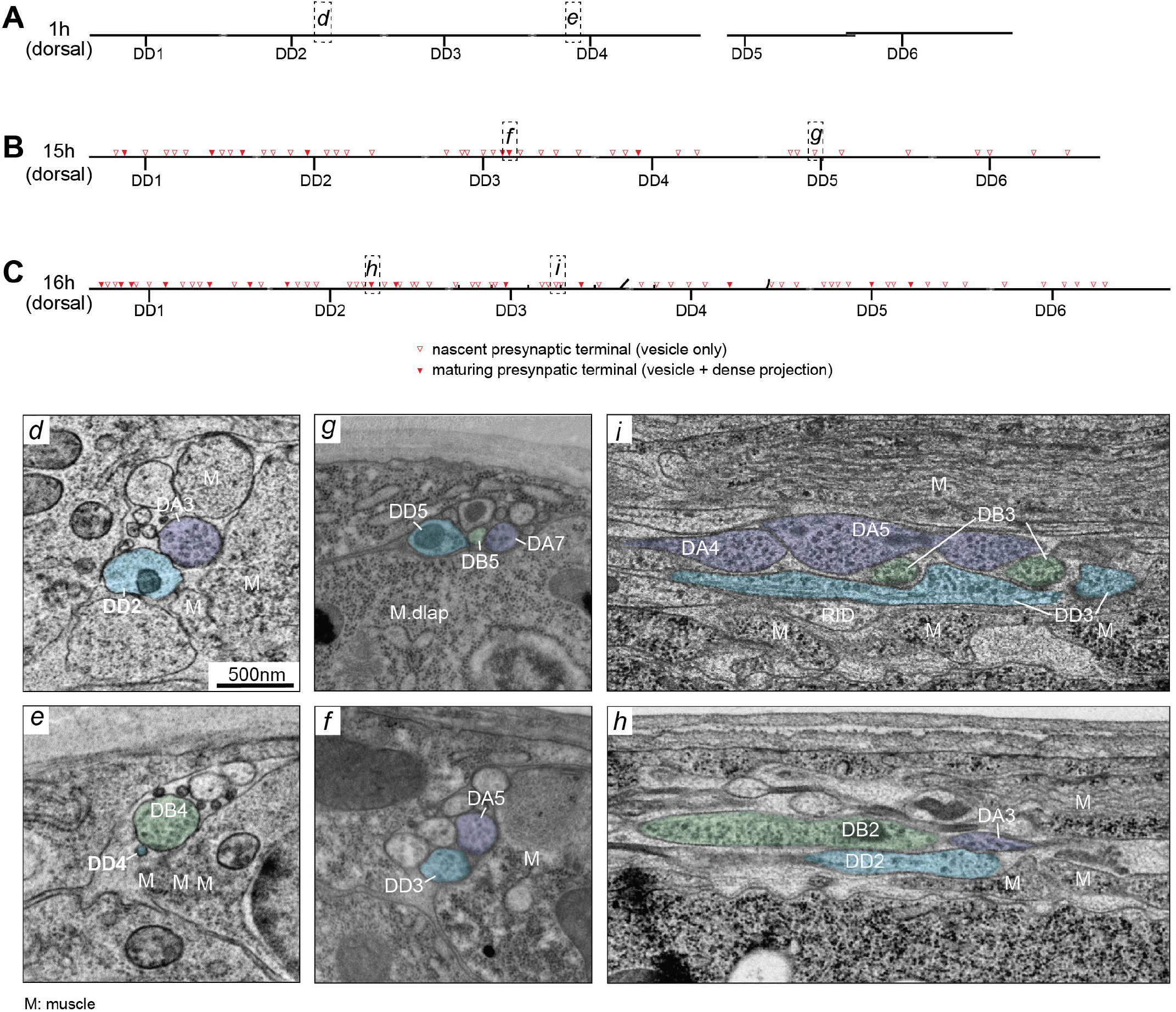
Examples of electron micrographs of DD dorsal neurite’s sequential development of NMJs. (A) At early L1, DD’s dorsal neurite is a dendrite, strictly post-synaptic to the embryonic DA (d) and DB (e) eMNs. (B-C) At the late L1 and L1/L2 transition stages, DD’s dorsal process has acquired axonal morphology, with developing presynaptic terminals, some with vesicles and presynaptic dense projections (f, h) and some with only vesicles (g, i).

**Figure 5 – Supp. 1.**
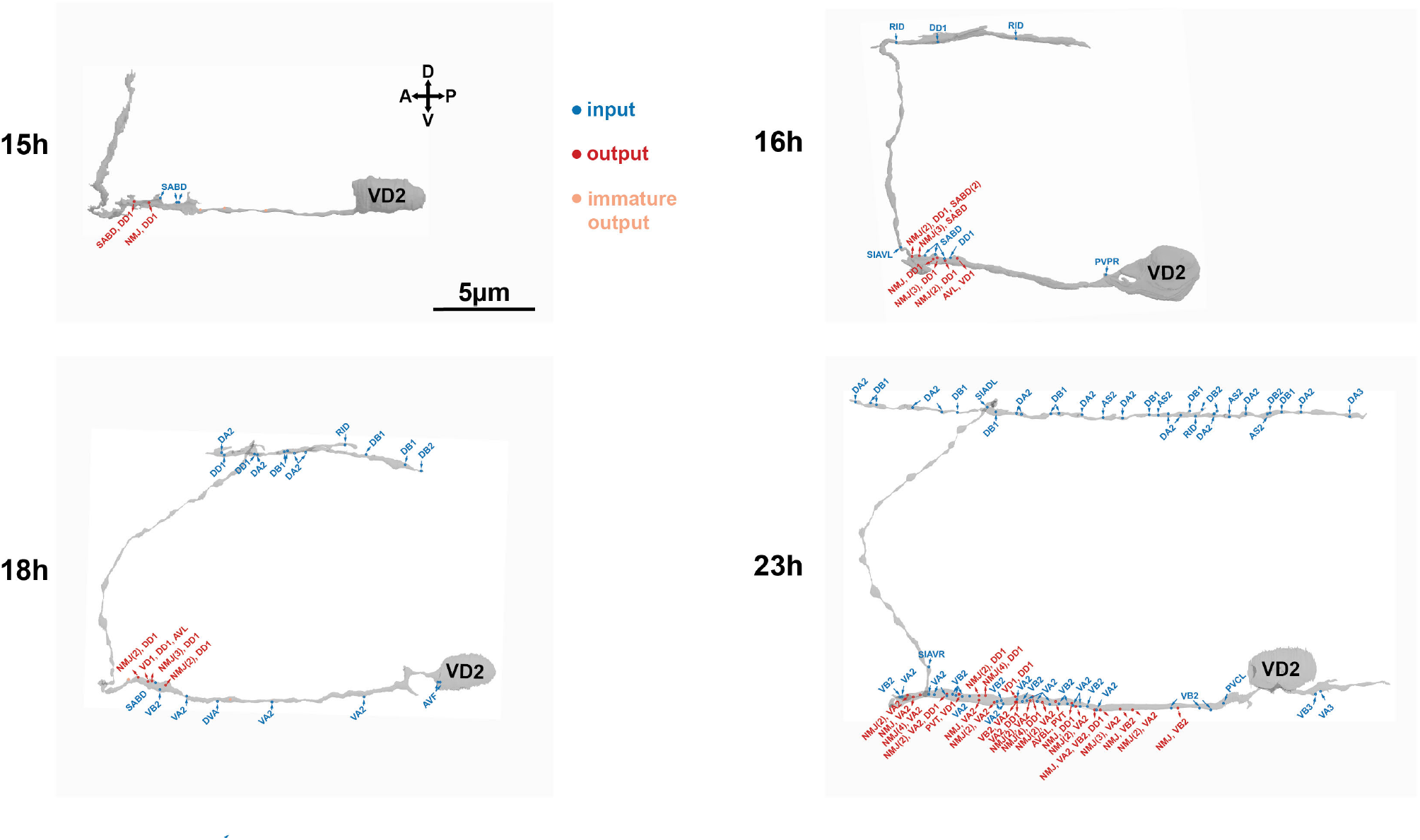
The establishment of connectivity in VD2 as it develops. Developmental wiring diagram of VD2 at early L1 (hr), late L1 (15hr), L1/L2 transition (16hr), early L2 (18hr) and late L2 (23hr). NMJ: neuromuscular junction. Numbers in brackets denote the number of chemical synapses for input or output cells. All other inputs and outputs have one chemical synapse.

**Figure 5 – Supp. 2.**
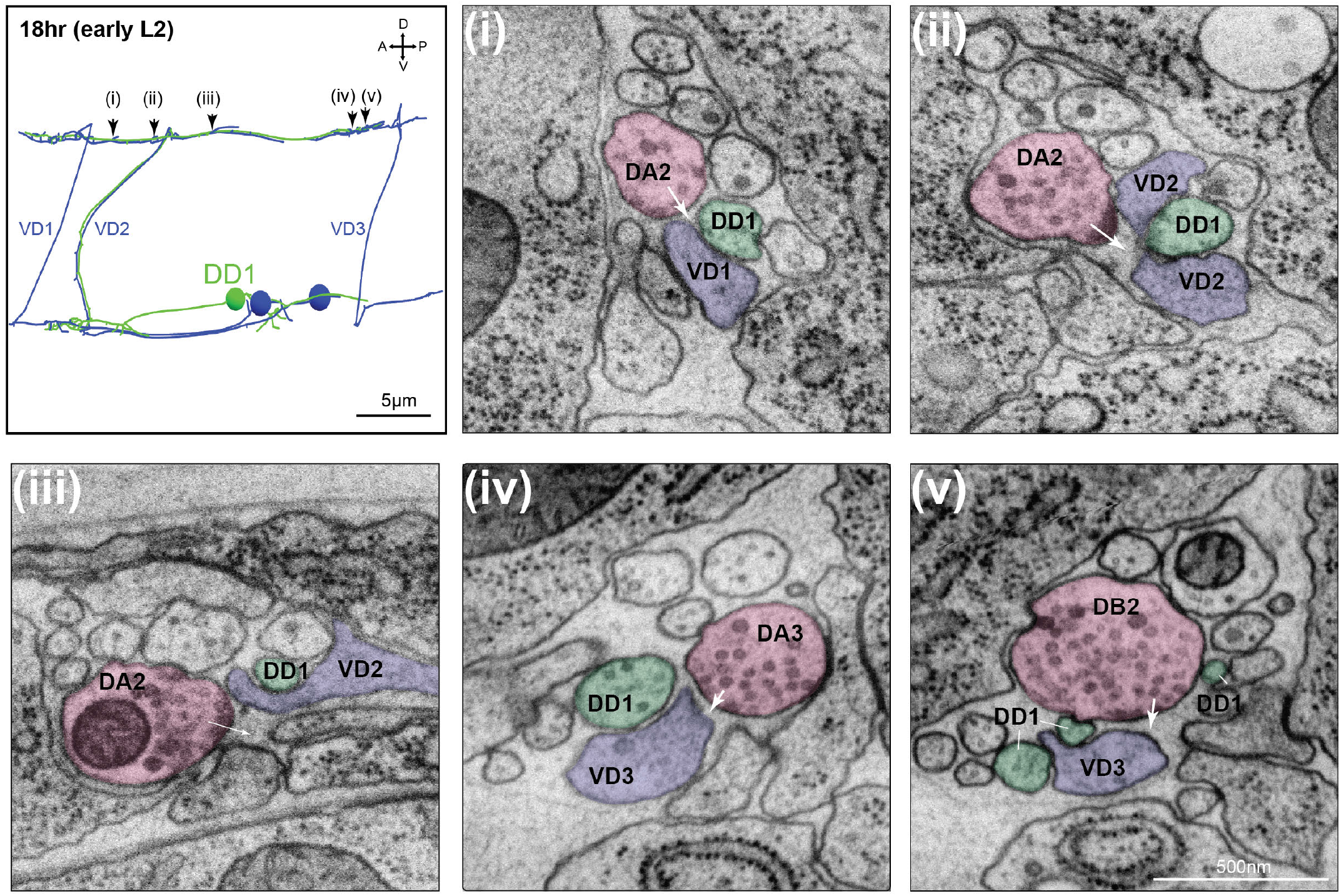
Examples of extending VD neurites entering the postsynaptic fields of embryonic eMN synapses. (Left panel) Skeleton reconstruction of neurites of DD1 and three post-embryonic VD at early L2 stage. i-V: Along the dorsal nerve cord, DD1 maintains its post-synaptic apposition to presynaptic terminals of embryonic DA (i-iv) and DB (v) eMNs, all with the growing VD neurites running adjacently. White arrows denote the post-synaptic fields of the respective presynaptic terminals.

**Figure 5 – Supp. 3.**
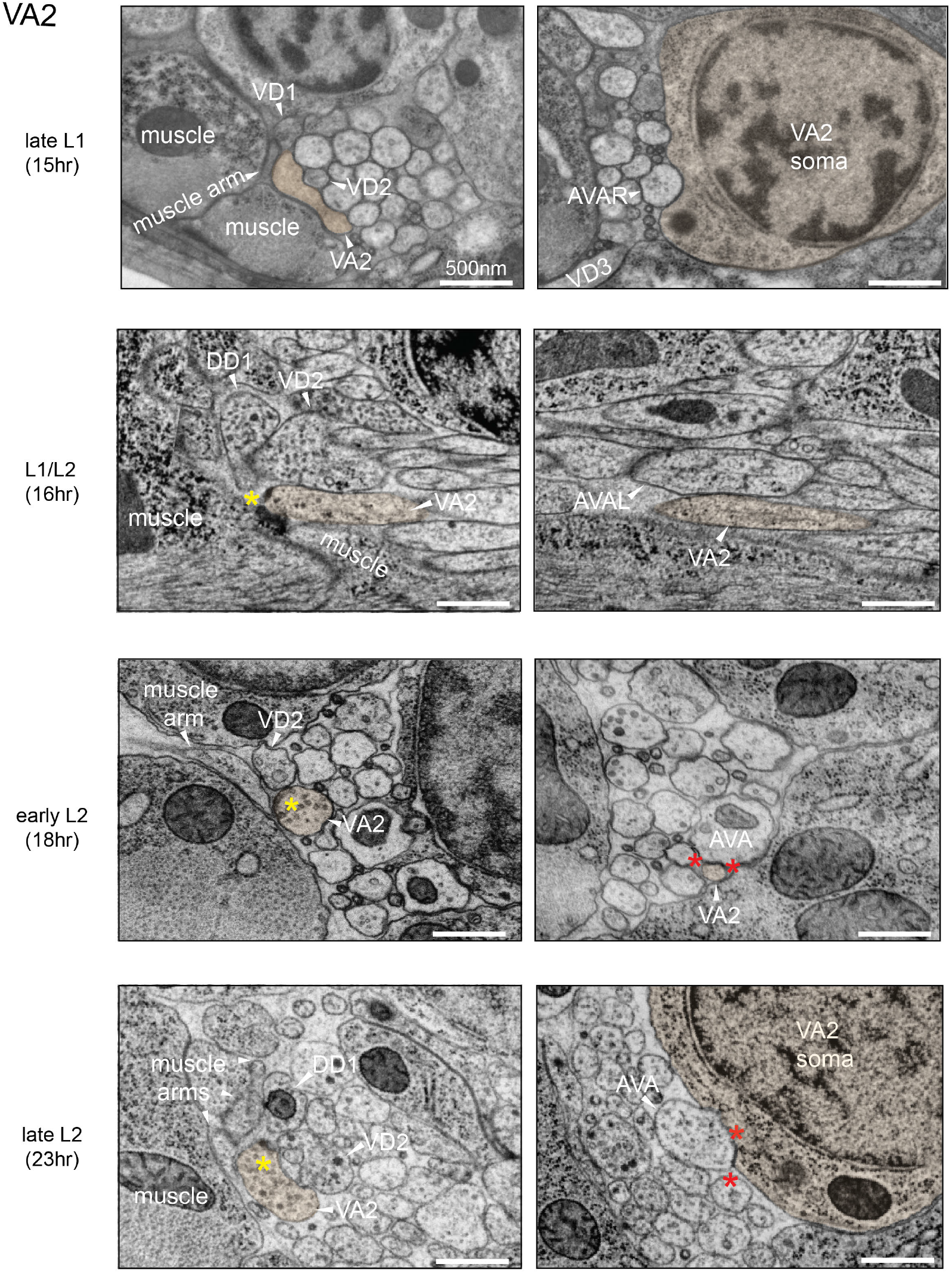
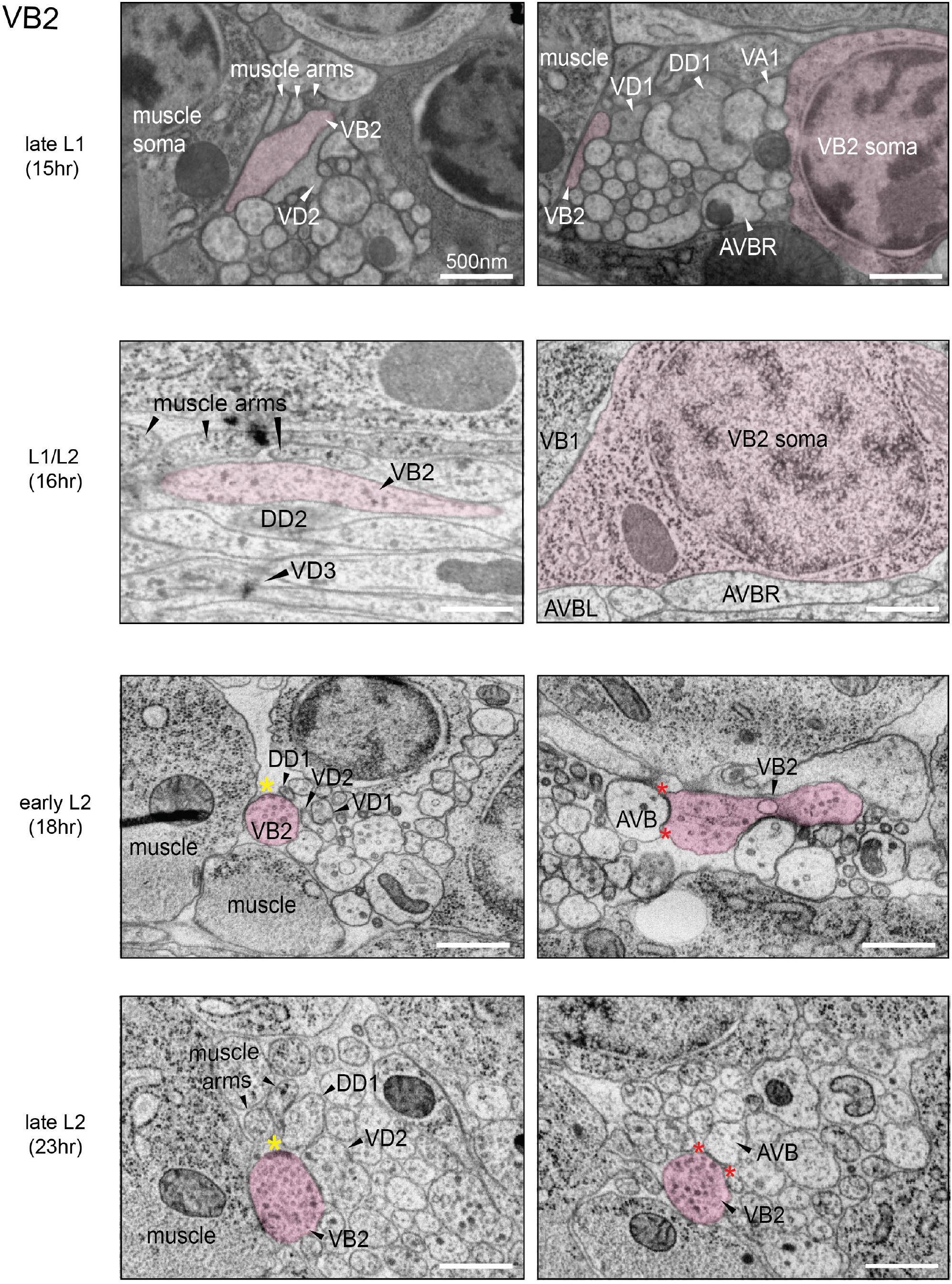
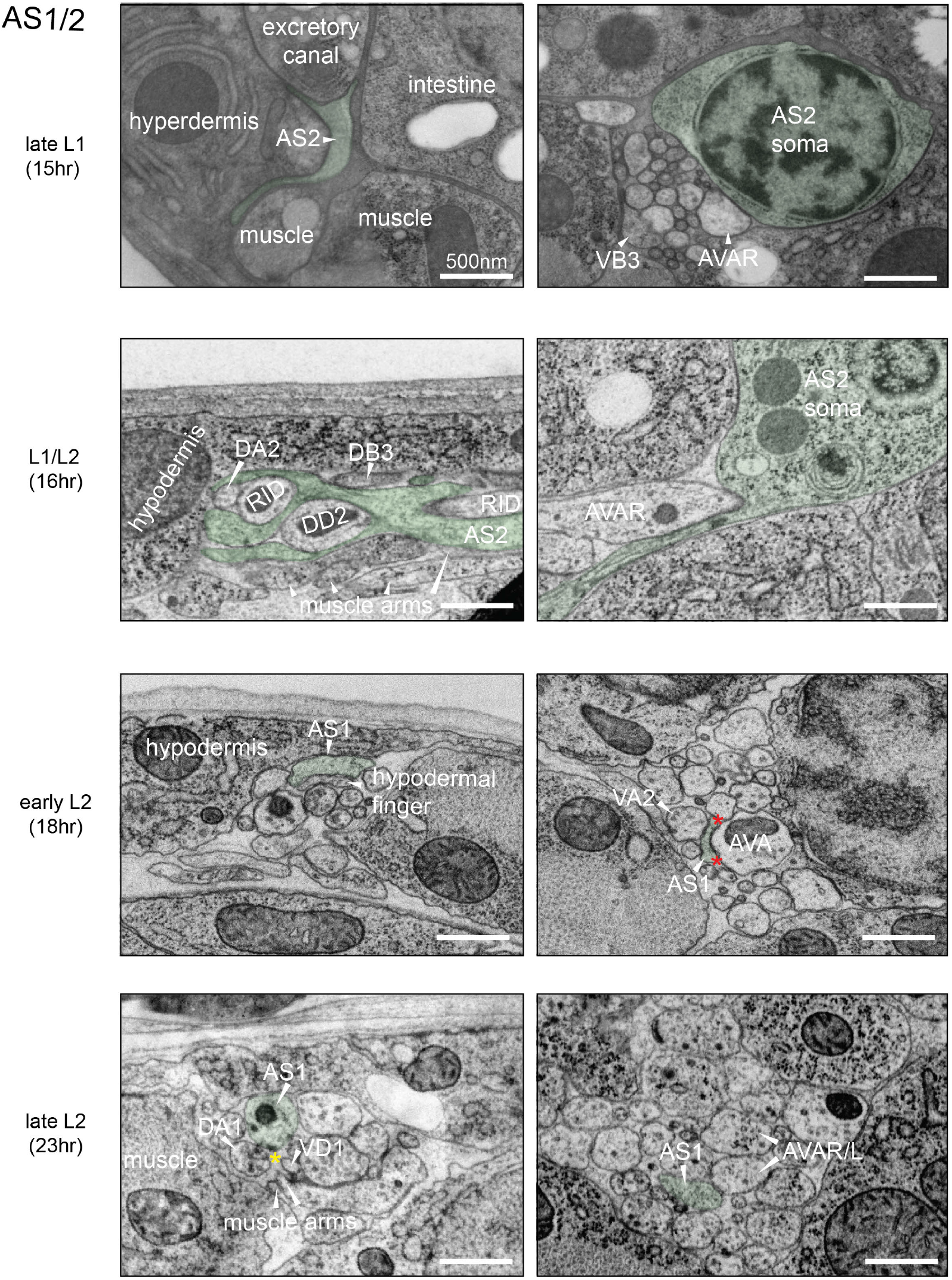
Post-embryonic eMN neurites exhibit programmed and sequential building of synapses. Example electron micrographs of the VA, VB and AS motor neurons at early L1 (1hr), late L1 (15hr), L1/L2 transition (16hr), early L2 (18hr), and late L2 (23hr). **(a) *VA2***: (Left panels:) VA2 neurite already elongates along muscles, in close proximity to the DD and VD neurites (1hr). As it migrates, it develops swellings in apposition to DD and muscles (its future post-synaptic partners), filling first with presynaptic dense projections (16hr), followed by vesicles (post 16hr). Yellow stars (*) denote maturing presynaptic terminals of VA2’s dyatic or polyatic NMJs. (Right panels:) VA2 neurite is already in contact with its future input partner, the premotor interneuron AVA, with putative gap junctions, bordered by red stars (*). **(b) *VB2***: (Left panels:) VB2’s migrating neurite already migrates along muscles (1hr). It develops presynaptic swellings in apposition to DD neurites and muscles, its future post-synaptic partners; some swellings accumulate vesicles (16hr) before adding presynaptic dense projections (post 16hr) in maturing, dyatic or polyatic NMJs, denoted by yellow stars (*). (Right panels:) By L1/L2 transition, it is already in contact with its future input partner premotor interneurons AVB with putative gap junctions, bordered by red stars (*). **(c) *AS***: (left panels:) AS1’s developing neurite migrates almost immediately towards the dorsal cord (1hr). In the dorsal cord, it wraps around DD and RID (16hr) while contacting muscle arms (16hr) and sometimes hypodermal fingers (18hr). It develops immature-looking presynaptic swellings in apposition to their future partners, muscles and VD (23hr). Developing NMJs denoted by yellow stars (*). (Right panels:) AS1’s short ventral process also makes contact and form putative gap junctions with the premotor interneurons AVA, bordered by red stars (*).

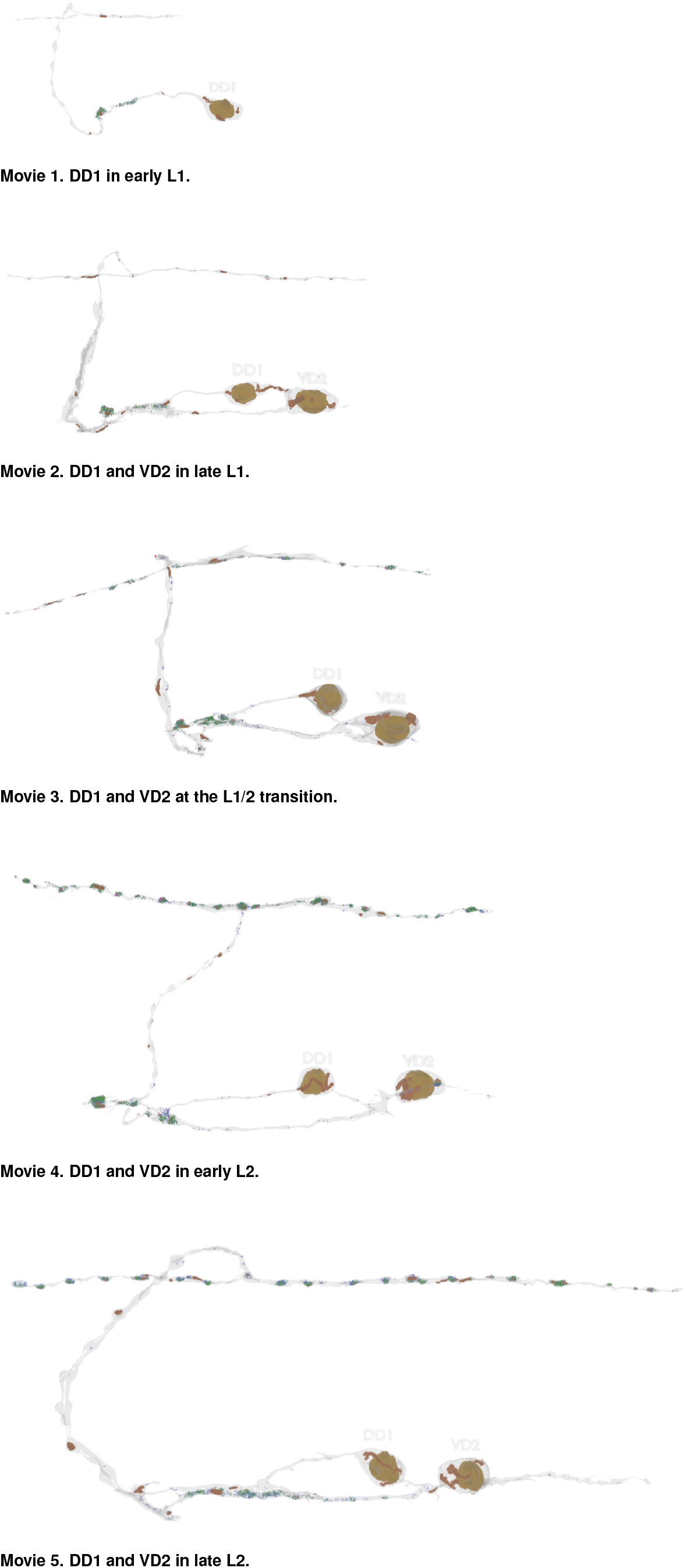

## Supplemental: Data sheets

**Data sheet 1. Commissural outgrowth by individual VD neuron at the late L1 stage**.

During commissural outgrowth, VD neuron interact with the epidermis (hypodermis), sublateral cords, and the excretory canal. Example electron micrographs and characteristic morphological features of individual VD neuron’s outgrowth are shown and described.

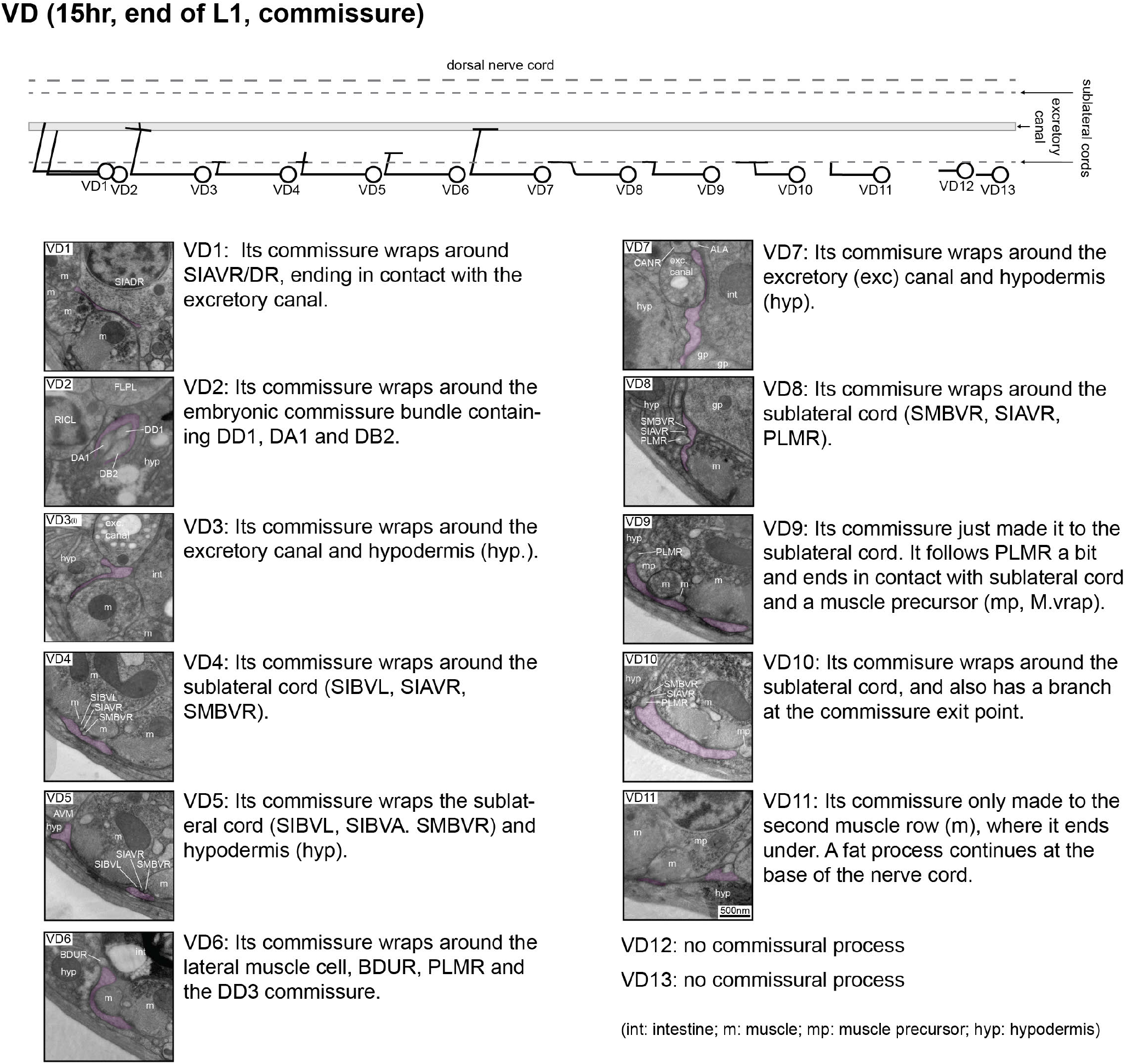

**Data sheet 2. Neurite outgrowth by individual VD neuron at the L1/L2 transition stage**.

In the ventral nerve cord, in addition to DD processes, many VD neurites make contacts with AVJ. During the later stages of VD commissural outgrowth, there is significant interaction with the excretatory canal, sublateral cords and epidermis (hypodermis), often engulfing chunks of epidermal tissues. Upon entering the dorsal nerve cord, VD processes swell up and fill space, and have a tendency to wrap the neurites of DD and RID. Example electron micrographs and characteristic morphological features of individual VD neuron’s outgrowth are shown and described.

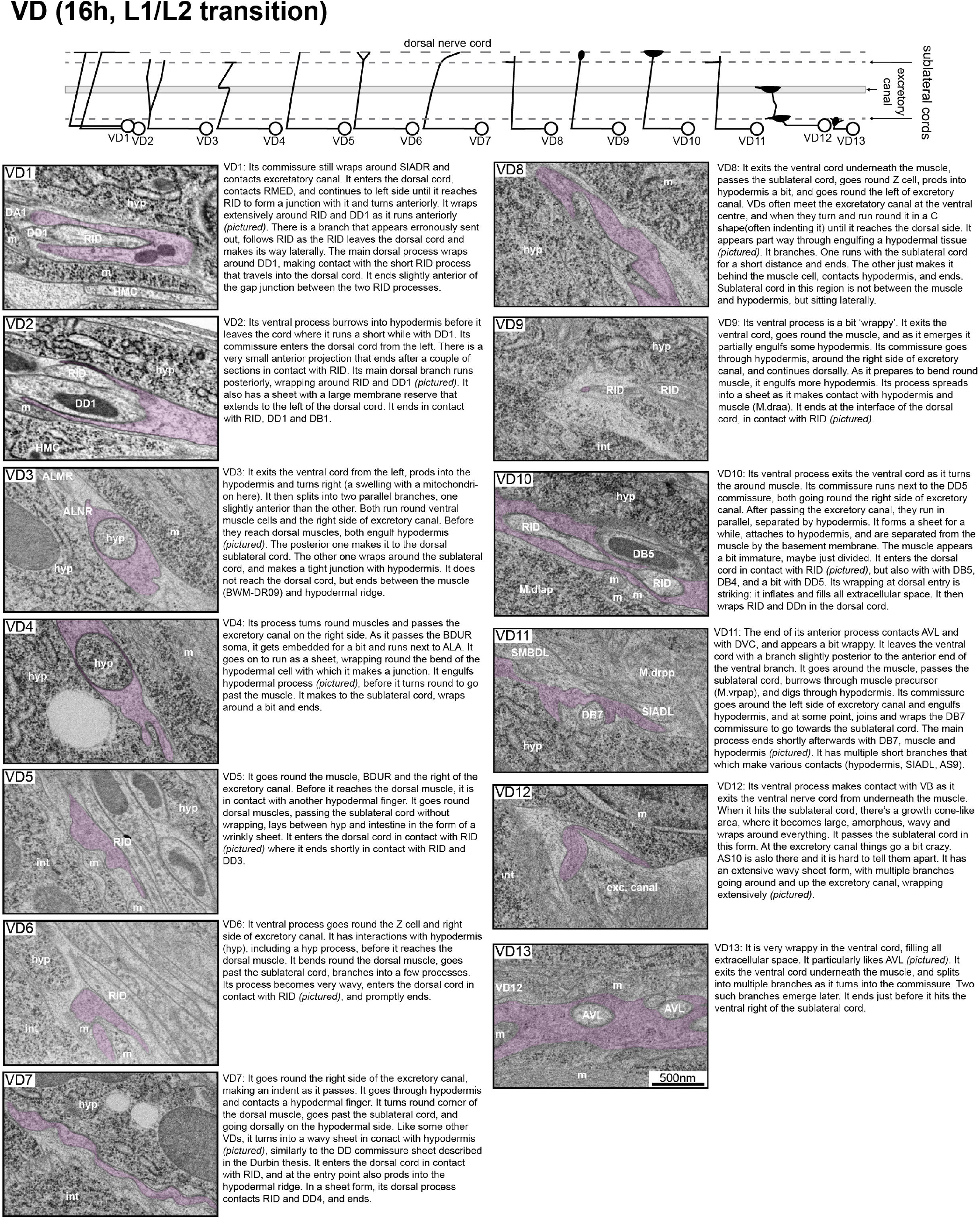

**Data sheet 3. Neurite outgrowth by individual AS neuron at the late L1 stage**.

During outgrowth in the ventral and dorsal nerve cords, as well as the circumferential commissural extension, AS neurites wrap extensively, frequently the sublateral cords, the excretory canal, and DD neurites. Their neurites dwell only briefly in the ventral cord, forming putative gap junctions with AVA, AVB and others neurons. Example electron micrographs and characteristic features of individual AS neuron outgrowth are shown and described.

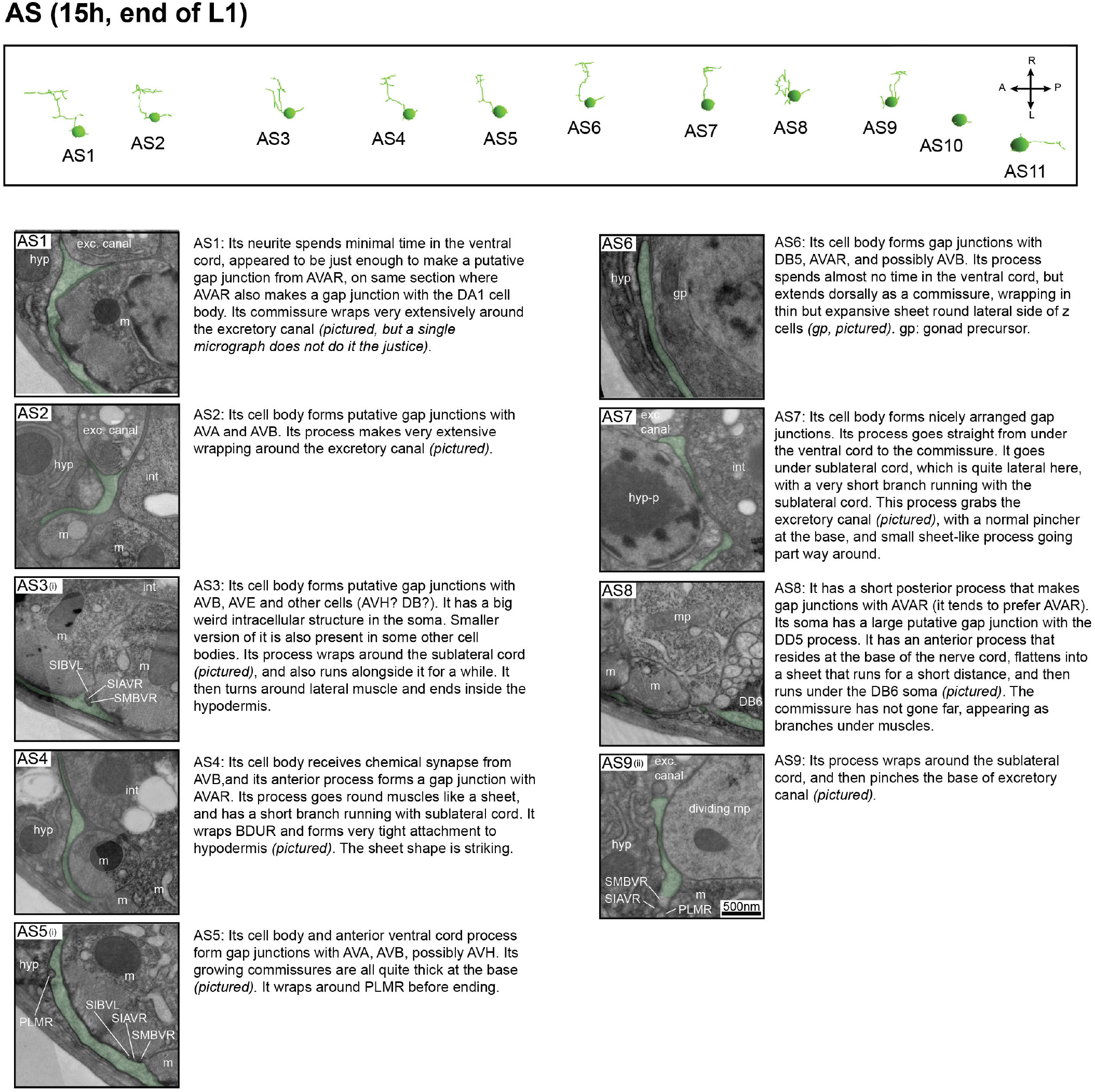

**Data sheet 4. Neurite outgrowth by individual VA neuron at the late L1 stage**.

VA neurites exhibit an immature, non-cylindrical morphology, sending sheet-like protrusions that are directed dorsally toward the area where NMJs are made by DD motor neurons in the ventral nerve cord. Compared to AS and VD, their posterior-to-anterior growth is quite unremarkable except that they often seem to contact PVC and likely DVA, and they make putative gap junctions with AVA. Example electron micrographs and characteristic features of individual VA neuron outgrowth are shown and described.)

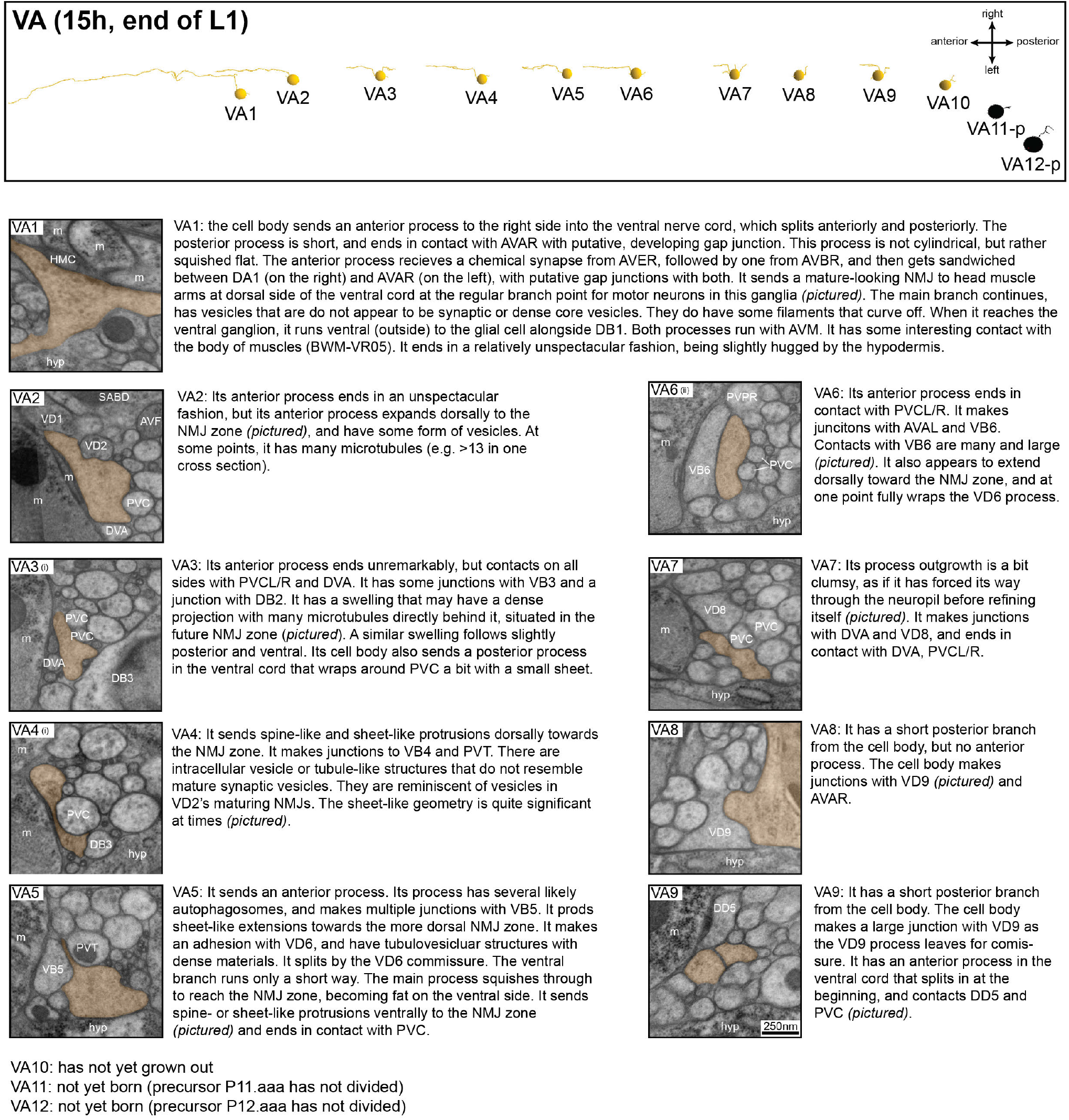

**Data sheet 5. Neurite outgrowth by individual VB neuron at the late L1 stage**.

Similar to the VA motor neurons, VB neurites adopt an immature, non-cylindrical morphology, sending sheet-like protrusions that are directed dorsally toward the area where NMJs are made by DD motor neurons in the ventral nerve cord. PVC and likely DVA make frequent contacts with multiple VB processes. Their soma and neurites make putative gap junctions with AVB. Example electron micrographs and characteristic features of individual AS neuron outgrowth are shown and described.)

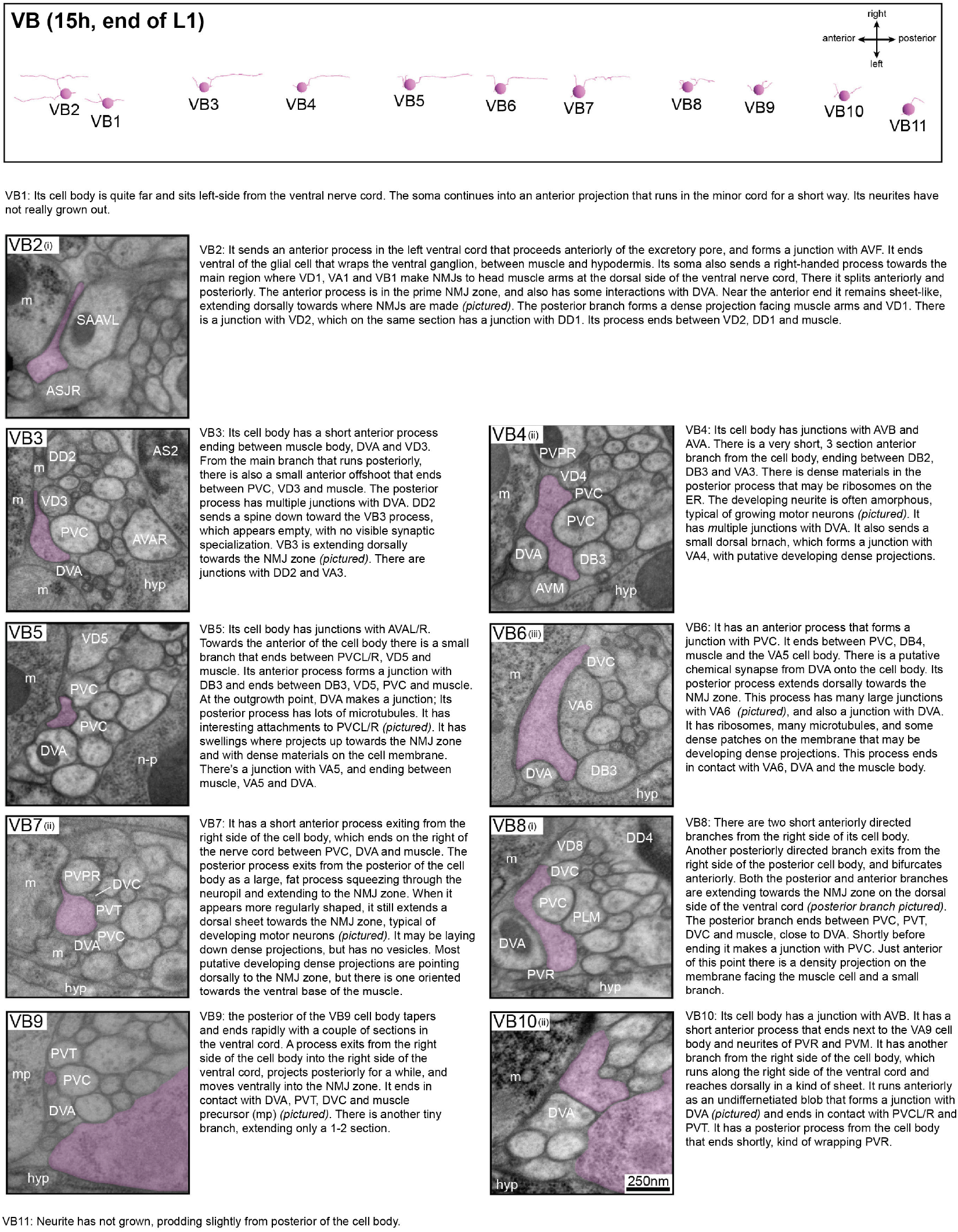

## References

[1] “Postnatal brain development”. In: The Cambridge Encyclopedia of Child Development. 2nd ed. Cambridge University Press, 2017, pp. 563–594. DOI: 10.1017/9781316216491.090.

[2] John E Sulston and H Robert Horvitz. “Post-embryonic cell lineages of the nematode, Caenorhabditis elegans”. In: Developmental biology 56.1 (1977), pp. 110–156.

[3] J.E. Sulston et al. “The embryonic cell lineage of the nematode Caenorhabditis elegans”. In: Developmental Biology 100.1 (1983), pp. 64–119. ISSN: 0012-1606. DOI: https://doi.org/10.1016/0012-1606(83)90201-4.

[4] John Edward Sulston and Sydney Brenner. “Post-embryonic development in the ventral cord of<i> Caenorhabditis elegans</i>“. In: Philosophical Transactions of the Royal Society of London. B, Biological Sciences 275.938 (1976), pp. 287–297. DOI: 10.1098/rstb.1976.0084.

[5] J. E. Sulston and H. R. Horvitz. “Post-embryonic cell lineages of the nematode, Caenorhabditis elegans”. In: Developmental biology 56.1 (1977), pp. 110–156.

[6] J.G. White, D.G. Albertson, and M.A. Anness. “Connectivity changes in a class of motoneurone during the development of a nematode”. In: Nature 271.5647 (1978), pp. 764–766.

[7] J. G. White et al. “The structure of the nervous system of the nematode Caenorhabditis elegans”. In: Philosophical Transactions of the Royal Society of London. Series B, Biological Sciences 314.1165 (1986), pp. 1–340. ISSN: 0962-8436.

[8] Steven J Hallam and Yishi Jin. “lin-14 regulates the timing of synaptic remodelling in Caenorhabditis elegans”. In: Nature 395.6697 (1998), pp. 78–82.

[9] John Graham White et al. “The structure of the ventral nerve cord of Caenorhabditis elegans</i>“. In: Philosophical Transactions of the Royal Society of London. B, Biological Sciences 275.938 (1976), pp. 327–348. DOI: 10.1098/rstb.1976.0086.

[10] Andrea Cuentas-Condori and David M Miller 3rd. “Synaptic remodeling, lessons from C. elegans”. In: Journal of Neurogenetics 34.3-4 (2020), pp. 307–322.

[11] Naina Kurup and Yishi Jin. “Neural circuit rewiring: insights from DD synapse remodeling”. In: Worm 5.1 (2016), e1129486. DOI: 10.1080/21624054.2015.1129486.

[12] Stephen E. Von Stetina, Millet Treinin, and David M. Miller. “The Motor Circuit”. In: The Neurobiology of C. elegans. Vol. 69. International Review of Neurobiology. Academic Press, 2005, pp. 125–167. DOI: https://doi.org/10.1016/S0074-7742(05)69005-8.

[13] K. Howell, J.G. White, and O. Hobert. “Spatiotemporal control of a novel synaptic organizer molecule”. In: Nature 523.7558 (2015), pp. 83–7. DOI: 0.1038/nature14545.

[14] Shay Stern, Christoph Kirst, and Cornelia I. Bargmann. “Neuromodulatory Control of Long-Term Behavioral Patterns and Individuality across Development”. In: Cell 171.7 (2018), 1648–1662.e10.

[15] Neil A Croll. “Components and patterns in the behaviour of the nematode Caenorhabditis elegans”. In: Journal of zoology 176.2 (1975), pp. 159–176.

[16] Yangning Lu et al. “Extrasynaptic signaling enables an asymmetric juvenile motor circuit to produce a symmetric gait”. In: bioRxiv (2021). DOI: 10.1101/2021.09.21.461278.

[17] Ramóny Cajal. “A quelle ceoque apparissent les expen-sions des cellules nerveuses de la molle epiniere du poulet.” In: Anat Anz erger 5 (1890), pp. 609–613.

[18] Ross Granville Harrison. “The outgrowth of the nerve fiber as a mode of protoplasmic movement”. In: Journal of Experimental Zoology 9.4 (1910), pp. 787–846. DOI: https://doi.org/10.1002/jez.1400090405. eprint: https://onlinelibrary.wiley.com/doi/pdf/10.1002/jez.1400090405. URL: https://onlinelibrary.wiley.com/doi/abs/10.1002/jez.1400090405.

[19] Richard Michael Durbin. Studies on the development and organisation of the nervous system of Caenorhabditis elegans. 1987.

[20] Kristen M. Harris and Richard J. Weinberg. “Ultrastructure of Synapses in the Mammalian Brain”. In: Cold Spring Harbor Perspectives in Biology 4.5 (2012). DOI: 10.1101/cshperspect.a005587.

[21] Daniel Witvliet et al. “Connectomes across development reveal principles of brain maturation”. In: Nature (2021), pp. 1–5.

[22] Bruce A Bamber et al. “The Caenorhabditis elegans unc-49 locus encodes multiple subunits of a heteromultimeric GABA receptor”. In: Journal of Neuroscience 19.13 (1999), pp. 5348–5359.

[23] Bruce A Bamber, Roy E Twyman, and Erik M Jorgensen. “Pharmacological characterization of the homomeric and heteromeric UNC-49 GABA receptors in C. elegans”. In: British journal of pharmacology 138.5 (2003), pp. 883–893.

[24] Shangbang Gao and Mei Zhen. “Action potentials drive body wall muscle contractions in Caenorhabditis elegans”. In: Proceedings of the National Academy of Sciences 108.6 (2011), pp. 2557–2562.

[25] Ping Liu, Bojun Chen, and Zhao-Wen Wang. “Postsynaptic current bursts instruct action potential firing at a graded synapse”. In: Nature communications 4.1 (2013), pp. 1–12.

[26] Manuela DAlessandro et al. “CRELD1 is an evolutionarily-conserved maturational enhancer of ionotropic acetylcholine receptors”. In: eLife 7 (2018), e39649. DOI: 10.7554/eLife.39649.

[27] Christopher Fang-Yen et al. “Biomechanical analysis of gait adaptation in the nematode Caenorhabditis elegans”. In: Proceedings of the National Academy of Sciences 107.47 (2010), pp. 20323–20328.

[28] Michael L Nonet. “Visualization of synaptic specializations in live C. elegans with synaptic vesicle protein-GFP fusions”. In: Journal of neuroscience methods 89.1 (1999), pp. 33–40.

[29] Christelle Gally and Jean-Louis Bessereau. “GABA Is Dispensable for the Formation of Junctional GABA Receptor Clusters in Caenorhabditis elegans”. In: Journal of Neuroscience 23.7 (2003), pp. 2591–2599. DOI: 10.1523/JNEUROSCI.23-07-02591.2003.

[30] Robby M. Weimer. “Preservation of C. elegans Tissue Via High-Pressure Freezing and Freeze-Substitution for Ultrastructural Analysis and Immunocytochemistry”. In: C. elegans: Methods and Applications. Ed. by Kevin Strange. Humana Press, 2006, pp. 203–221. ISBN: 978-1-59745-151-2. DOI: 10.1385/1-59745-151-7:203.

[31] Ben Mulcahy et al. “A pipeline for volume electron microscopy of the Caenorhabditis elegans nervous system”. In: Frontiers in Neural Circuits 12 (2018), p. 94. ISSN: 1662-5110. DOI: 10.3389/fncir.2018.00094.

[32] Justin Gage Crump et al. “The SAD-1 kinase regulates presynaptic vesicle clustering and axon termination”. In: Neuron 29.1 (2001), pp. 115–129.

[33] Joanne SM Kim et al. “A chemical-genetic strategy reveals distinct temporal requirements for SAD-1 kinase in neuronal polarization and synapse formation”. In: Neural development 3.1 (2008), pp. 1–14.

[34] Wesley Hung et al. “Neuronal polarity is regulated by a direct interaction between a scaffolding protein, Neurabin, and a presynaptic SAD-1 kinase in Caenorhabditis elegans”. In: Development 134.2 (2007), pp. 237–249.

[35] Mikyoung Park et al. “CYY-1/cyclin Y and CDK-5 differentially regulate synapse elimination and formation for rewiring neural circuits”. In: Neuron 70.4 (2011), pp. 742–757.

[36] Naina Kurup et al. “Dynamic microtubules drive circuit rewiring in the absence of neurite remodeling”. In: Current Biology 25.12 (2015), pp. 1594–1605.

[37] Jun Meng et al. “Myrf ER-bound transcription factors drive C. elegans synaptic plasticity via cleavage-dependent nuclear translocation”. In: Developmental cell 41.2 (2017), pp. 180–194.

[38] WW Walthall and Jeffery A Plunkett. “Genetic transformation of the synaptic pattern of a motoneuron class in Caenorhabditis elegans”. In: Journal of Neuroscience 15.2 (1995), pp. 1035–1043.

[39] Jerome Korzelius et al. “C. elegans MCM-4 is a general DNA replication and checkpoint component with an epidermis-specific requirement for growth and viability”. In: Developmental biology 350.2 (2011), pp. 358–369.

[40] WW Walthall et al. “Changing synaptic specificities in the nervous system of Caenorhabditis elegans: differentiation of the DD motoneurons”. In: Journal of neurobiology 24.12 (1993), pp. 1589–1599.

[41] Yishi Jin, Roger Hoskins, and H Robert Horvitz. “Control of type-D GABAergic neuron differentiation by C. elegans UNC-30 homeodomain protein”. In: Nature 372.6508 (1994), pp. 780–783.

[42] Philbrook Alison et al. “Neurexin directs partner-specific synaptic connectivity in C. elegans”. In: eLife 7 (2018).

[43] Katherine L Thompson-Peer et al. “HBL-1 patterns synaptic remodeling in C. elegans”. In: Neuron 73.3 (2012), pp. 453–465.

[44] Tyne W Miller-Fleming et al. “The DEG/ENaC cation channel protein UNC-8 drives activity-dependent synapse removal in remodeling GABAergic neurons”. In: Elife 5 (2016), e14599.

[45] Tyne W Miller-Fleming et al. “Transcriptional control of parallel-acting pathways that remove specific presynaptic proteins in remodeling neurons”. In: Journal of Neuroscience 41.27 (2021), pp. 5849–5866.

[46] Ami Citri and Robert C Malenka. “Synaptic plasticity: Multiple forms, functions, and mechanisms”. In: Nature Neuropsychopharmacology 33 (2007), pp. 18–41.

[47] S. J. Martin, P. D. Grimwood, and R. G. M. Morris. “Synaptic Plasticity and Memory: An Evaluation of the Hypothesis”. In: Annual Review of Neuroscience 23.1 (2000), pp. 649–711. DOI: 10.1146/annurev.neuro.23.1.649.

[48] Cornelia I Bargmann and Eve Marder. “From the connectome to brain function”. In: Nature Methods 10.6 (2013), pp. 483–490. DOI: 10.1038/nmeth.2451.

[49] Victoria J. Butler et al. “A consistent muscle activation strategy underlies crawling and swimming in Caenorhabditis elegans”. In: Journal of The Royal Society Interface 12.102 (2015), p. 20140963. DOI: 10.1098/rsif.2014.0963.

[50] Arthur D Edelstein et al. “Advanced methods of microscope control using μManager software”. In: Journal of Biological Methods 1.2 (2014), e10. DOI: 10.14440/jbm.2014.36. URL: https://jbmethods.org/jbm/article/view/36.

[51] Taizo Kawano et al. “An Imbalancing Act: Gap Junctions Reduce the Backward Motor Circuit Activity to Bias C. elegans for Forward Locomotion”. In: Neuron 72.4 (2011), pp. 572–586. DOI: https://doi.org/10.1016/j.neuron.2011.09.005.

[52] R. Schalek et al. “ATUM-based SEM for High-Speed Large-Volume Biological Reconstructions”. In: Microscopy and Microanalysis 18.S2 (2012), pp. 572–573. DOI: 10.1017/S1431927612004710.

[53] Kenneth J. Hayworth et al. “Imaging ATUM ultrathin section libraries with WaferMapper: a multi-scale approach to EM reconstruction of neural circuits”. In: Frontiers in Neural Circuits 8 (2014), p. 68. DOI: 10.3389/fncir.2014.00068.

[54] Johannes Schindelin et al. “Fiji: an open-source platform for biological-image analysis”. In: Nat Methods 9 (2012), pp. 676–682. DOI: doi.org/10.1038/nmeth.2019.

[55] Albert Cardona et al. “TrakEM2 Software for Neural Circuit Reconstruction”. In: PLOS ONE 7.6 (2012), pp. 1–8. DOI: 10.1371/journal.pone.0038011.

[56] Stephan Saalfeld et al. “CATMAID: collaborative annotation toolkit for massive amounts of image data”. In: Bioinformatics 25.15 (2009), pp. 1984–1986. DOI: 10.1093/bioinformatics/btp266.

[57] Daniel R. Berger, H. Sebastian Seung, and Jeff W. Lichtman. “VAST (Volume Annotation and Segmentation Tool): Efficient Manual and Semi-Automatic Labeling of Large 3D Image Stacks”. In: Frontiers in Neural Circuits 12 (2018), p. 88. DOI: 10.3389/fncir.2018.00088.

[58] Giusto J Ruvkun G. “The Caenorhabditis elegans heterochronic gene lin-14 encodes a nuclear protein that forms a temporal developmental switch”. In: Nature 338.6213 (1989), pp. 313–9. DOI: 10.1038/338313a0.

[59] David M. Miller and Diane C. Shakes. “Chapter 16 Immunofluorescence Microscopy”. In: Cuenorhubditis elegans: Modern Biologcal Analysis of an Organism. Ed. by Henry F. Epstein and Diane C. Shakes. Vol. 48. Methods in Cell Biology. Academic Press, 1995, pp. 365–394. DOI: https://doi.org/10.1016/S0091-679X(08)61396-5.

